# Rapidly Reconfigurable Dynamic Computing in Neural Networks with Fixed Synaptic Connectivity

**DOI:** 10.1101/2025.10.05.680523

**Authors:** Kai Mason, Sonia Sennik, Claudia Clopath, Aaron Gruber, Wilten Nicola

**Affiliations:** Department of Cell Biology and Anatomy, Cumming School of Medicine, University of Calgary, Calgary, Alberta, Canada; Creative Destruction Lab; Department of Bioengineering, Faculty of Engineering, Imperial College London, London, United Kingdom

## Abstract

Learning and memory in the brain’s neocortex have long been hypothesised to be primarily mediated by synaptic plasticity. Extensive research in artificial neural networks has shown that training networks by adjusting connection weights faces computational challenges, including large parameter spaces and the tendency of new learning to interfere with previous learning (catastrophic forgetting). We propose that the brain, which is resistant to these challenges, can also learn by modulating the excitability of each neuron in a network rather than changing synaptic strengths. We show here that learning a task-specific set of bias currents enables a feedforward or recurrent network with fixed and randomly assigned connections to perform well on and switch between dozens of tasks, including regression, classification, autonomous time series generation, a game and robotic control. Bias-only learning also provides a novel mechanistic explanation for representational drift. It directly links the noise robustness of neuronal representations on short and long time scales to the ability of neural circuits to preserve learned information while remaining adaptable. We postulate that subcortical structures, such as the basal ganglia or cerebellum, may provide similar bias inputs to the neocortex for rapid task learning and robustness against interference.

## Introduction

The adjustment of synaptic interconnections among neocortical neurons is widely postulated to be a key process in the formation of memories and task learning in the mammalian brain [1, 2]. It is unlikely, however, that such synaptic plasticity is the only mechanism of information storage or processing in the brain. In parallel series of developments in machine learning, increasingly complex network architectures with large parameter spaces are routinely suggested and explored as avenues of research [3], with some recent work focusing on modifying neuronal operations rather than network connectivity [4].

The modulation of cortical inputs by upstream structures [5, 6, 7, 8] and intrinsic changes in neuronal excitability [9, 10, 11] are alternate modes of information processing. In the context of motor learning and control, *in vivo* experimental evidence from the primary motor and dorsal premotor cortices of macaques has shown invariance in the local functional connectivity and neural covariance during motor adaptation tasks [7]. A computational modelling study of the same task utilising recurrent neural networks concluded that upstream adaptations were consistent with observed experimental data [8], which was also consistent with an earlier study in humans [12]. Similar results suggest that the encoding of visual, taste and auditory sensory information can be modulated by subcortical projections to the cortex [13, 14, 15]. These results suggest that adaptive processing in the neocortex may not require local changes in synaptic connectivity but may arise from changes in neuronal excitability or changes in upstream currents.

The computational utility of neuronal excitability adaptation remains to be fully explored. Recent studies in machine learning have begun researching how modification of activation functions affects network performance, such as fine-tuning after learning weights [16]. Other recent work has shown that networks with fixed random weights can perform some tasks, such as control or classification [17] by modifying so-called bias-currents, which mimic changes to neuronal excitability in computational models. However, these approaches fall short on the kinds of problems where cortical circuits appear to excel—for example, learning to approximate complex dynamical controllers. It is therefore not clear from prior work that changes to neuronal excitability via bias currents alone are sufficient to account for cortical information processing.

We consider here the extreme case in which neuronal connectivity is random and fixed, and test whether adjusting a neuronal excitability alone is sufficient to perform a variety of common dynamical tasks using low-rank recurrent neural networks [18, 19, 20, 21, 22]. In this paradigm, each neuron receives a bias current which can subtly shift its excitability. The bias currents represent input from subcortical or cortical structures. We henceforth refer to this as the bias adaptive neural firing framework (BANFF). We test here if recurrent BANFF networks can learn a broad range of more complex tasks, including learning closed-loop dynamical systems and autonomously producing the attractors of these systems. We found that BANFF networks could readily produce complex low-dimensional dynamics and switch between these dynamics instantly. Furthermore, bias-only training in BANFF RNNs provides a mechanistic explanation for both representational drift and robust computation with noisy units: networks can learn and switch between multiple, non-unique bias solutions to the same task on short (millisecond) and long (days) time scales. Further, bias-only learning in feed-forward architectures performed well in a variety of tasks using only one network. Importantly, the reduced parameter dimensionality (typically an order of magnitude smaller than the full weight set) affords practical advantages for training efficiency and hardware implementation in neuromorphic and real-time learning systems. This study supports the hypothesis that changes to neural excitability alone are sufficient for universal and robust computations.

## Results

### Learning with Bias Currents

Biologically, bias currents can be interpreted as presynaptic inputs, neuromodulators, or even changes to ion channel densities or types. However, the physiological effect on a neuron is to make the neuron more or less excitable (Figure 1A) by shifting its frequency-current relationship (*f* (*I*)-curve). The *f* (*I*)-curve classically measures the rate of spikes fired by a given input current, with more excitable neurons firing more spikes for a given input current (Figure 1B) [23].

**Figure 1.**
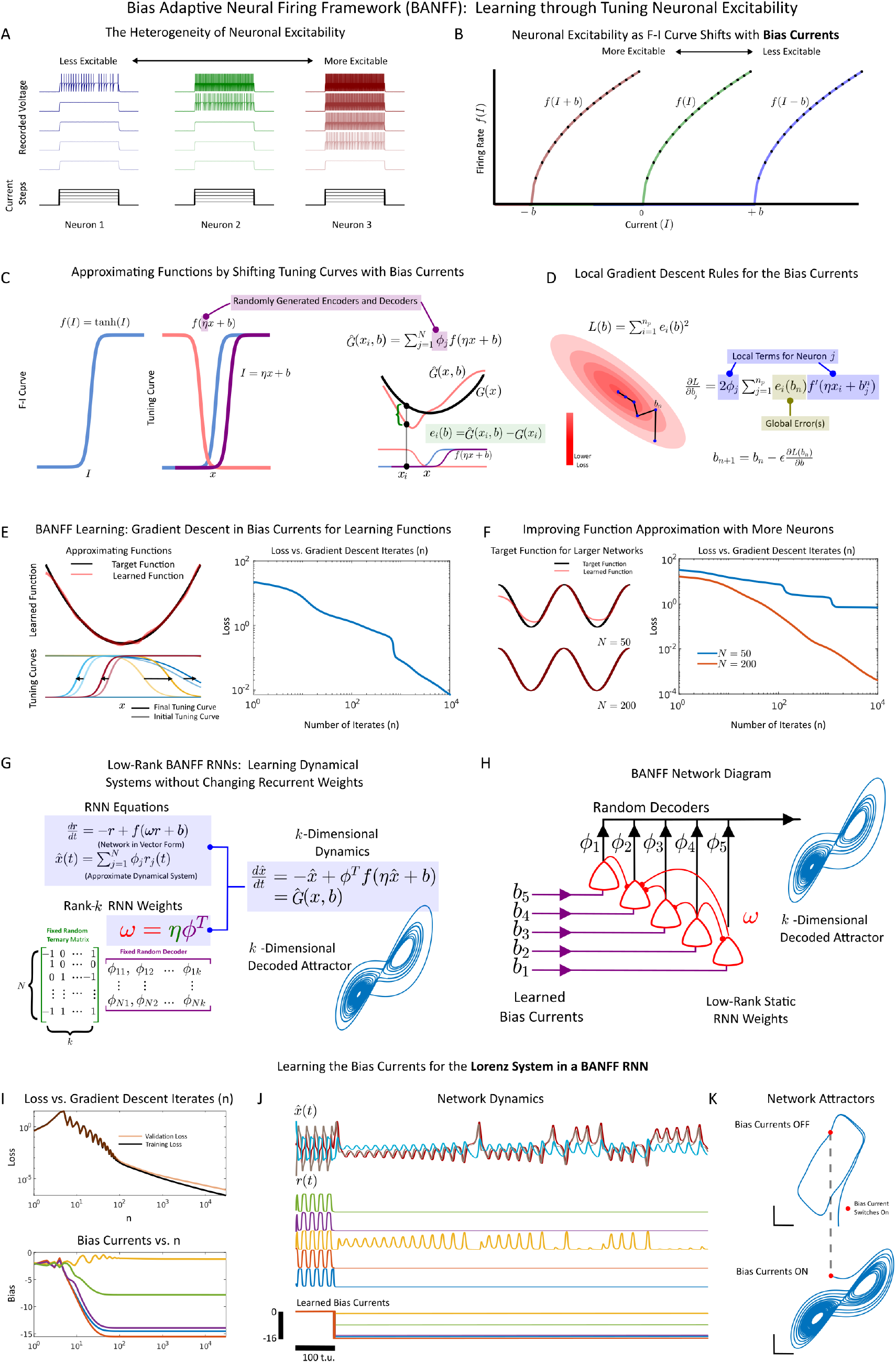
The Bias Adaptive Neural Firing Frame (BANFF): Learning Through Changes to Neural Excitability. **(A)** Illustration of heterogeneous responses to fixed applied step-currents (bottom). More excitable neurons fire at a higher spike rate for a given input current, while less excitable ones fire at lower rates. **(B)** The heterogeneity of neuronal excitability can be parameterised by a single variable: the bias current. The bias current shifts the *f* (*I*) curve, defined as the steady state firing rate as a function of the current *I*. More excitable neurons have *f* (*I*) curves shifted further to the right. **(C)** (Left) In recurrent neural networks, the closest analogue to the neuronal *f* (*I*) curve is the transfer function, which can be taken as *f* (*I*) = tanh(*I*). (Right) The input that a neuron receives is linearly expanded as *I* = *ηx* + *b*, where *b* acts as a bias current and *η* is the linear encoder for the variable *X*. Functions can be approximated by using a linear combination of tuning curves: 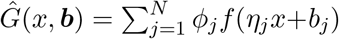. The encoders, *eta*_*j*_ and decoders *ϕ*_*j*_ are randomly generated and fixed. **(D)** Gradient descent can be used to determine the optimal bias currents with a local learning rule. Neuron *j*’s bias current is updated based on a global error signal, and the local properties of its own tuning curve *f*^*′*^(*η*_*j*_*x* + *b*_*j*_). **(E)** (Left) A network of *X* neurons (red) is trained to learn the function *y* = *x*^2^ (black) by only modifying the bias currents with local gradient descent. Each neuron has its initial bias current shifted during learning. (Right) The loss (mean-squared-error) versus the number of learning iterations. **(F)** (Left) A network of 50 neurons (top) and a larger network of 200 neurons (bottom) approximating the function *y* = *cos*(*πx*) on *x ∈* [*−*1, 1] through gradient descent in the bias currents. As more neurons are used, the network is better able to approximate *y* = *cos*(*πx*). (Right) The loss versus the number of iterations. **(G)** A low-rank BANFF recurrent neural network. The RNN is governed by the dynamical system highlighted in blue, with the decoded dynamics given by 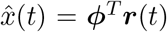. The weights for the RNN are determined by ***ω*** = ***ηϕ***^*T*^, and the encoders and decoders are randomly generated. By using a low-rank weight matrix, the RNN collapses onto the low-dimensional dynamical system given 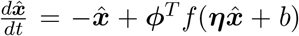. **(H)** For low-rank BANFF-RNNs, determining the correct set of bias currents through gradient descent will allow the RNN to act as a universal approximator for dynamical systems. **I** (Top) A low-rank BANFF RNN was trained to mimic the Lorenz system dynamics. The training (black) and validation (orange) loss is shown. (Bottom) The evolution of the bias currents for 5 neurons through gradient descent. **J** A simulation of the BANFF RNN where the bias currents are switched on after 100 time units (arbitrary units). The dynamics of 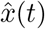 (top) versus the dynamics of 5 neurons (bottom). **K** The decoded attractor for the BANFF RNN with the bias currents off (top) and on (bottom). The BANFF RNN mimics the dynamics of the Lorenz system.

To model bias-only learning with the Bias-Adaptive-Neural Firing Framework (BANFF), we first considered recurrent neural networks due to their widespread use in modelling time-dependent processing in biological circuits [24, 25]:

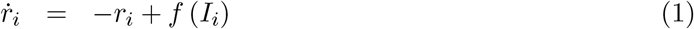

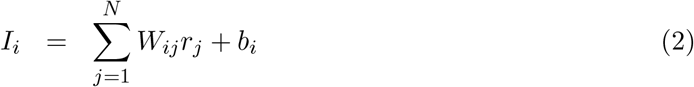

where *f* (*I*) is the neuron transfer function, *b*_*i*_ is the bias, and *W*_*ij*_ is a randomly generated weight that couples neuron *j* to neuron *i*. The variable *I*_*i*_ is the total current for neuron *i* while the variable *r*_*i*_(*t*) is the filtered firing rate for neuron *i*. The bias current shifts the transfer function *f* (*I*) of a neuron to more (*b >* 0) or less (*b <* 0) excitable regimes. In this work, we used *f* (*I*) = tanh(*I*) as the transfer function (Figure 1C), as it qualitatively captures the characteristics of neuronal *f* (*I*)-curves, namely saturation at very low and very high currents.

To first demonstrate how bias-only learning works in BANFF-trained networks, we considered low-rank RNNs, where

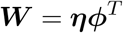

where ***η*** and ***ϕ*** are the neural encoder and decoder, respectively. The low-rank assumption provides critical insight into how RNNs/ANNs can learn by modifying bias currents only. Furthermore, the computational power of low-rank neural networks has been thoroughly explored in the Neural Engineering Framework [21, 22, 26] and other studies that assume random low-rank coupling [18, 19, 20].

By applying the low-rank assumption, the RNNs display low-dimensional dynamics where the dynamical variable of interest, 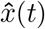, is governed by the differential equations

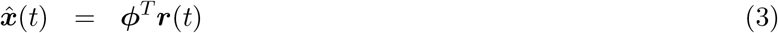

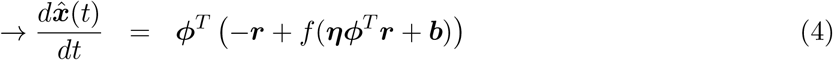

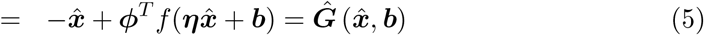

where the dynamics are given by the function 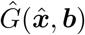, which is parameterized by the vector of bias currents ***b***. Thus, for the RNN to learn the dynamics of the system 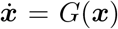, the bias currents ***b*** had to be learned to drive 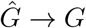 (Figure 1C). This was accomplished by minimising the squared loss:

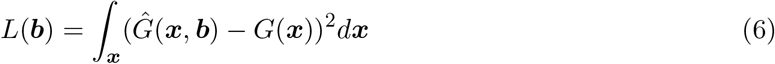

by computing the gradient with respect to the loss (Figure 1D), ∇*L*_***b***_ and updating ***b*** iteratively as follows:

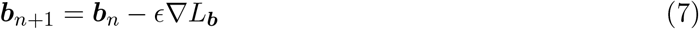

where *ϵ* controls the learning rate. Note that we use a momentum term to adjust the bias currents along with the gradient (Materials and Methods).

First, we note that the gradient only depends on a global error signal and local information specific to neuron *i* in low-rank BANFF networks (Figure 1D) as:

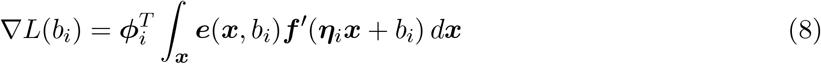

By applying this rule, any static function, *G*(***x***), can be learned iteratively with gradient descent operating exclusively on bias currents (Figure 1E). For example, the quadratic function *f* (*x*) = *x*^2^ was learned through BANFF learning (Figure 1E). As in traditional neural networks, where all weights are free parameters, adding more neurons improved network performance (Figure 1F).

By learning a static function with bias currents, the low-rank assumption allows a recurrent neural network to mimic the dynamics of 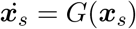 by learning the function *G*(***x***) with 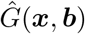 (Figure 1G). This allowed a randomly coupled low-rank network to learn low-dimensional dynamical systems (Figure 1H).

First, we tested BANFF-learning with the Lorenz system:

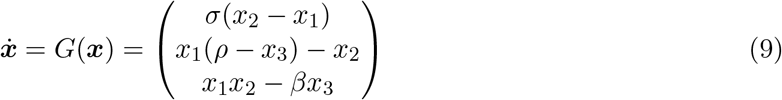

We found that the bias currents could be readily learned by using gradient descent with momentum (Figure 1I), with the bias currents converging to a steady state (Figure 1I). When tested, BANFF networks could readily produce the low-dimensional Lorenz butterfly attractor (Figure 1J-K) when the bias currents were applied. The network switched between its default attractor (a low-dimensional oscillator for ***b*** = 0) and the Lorenz system instantly.

We next investigated the generalisability of the Bias-Adaptive-Neural-Firing Framework by testing if a single BANFF network could learn multiple dynamical systems of different dimensions and subsequently switch between them (Figure 2A). We trained a single recurrent BANFF network with fixed and randomly generated weights (Figure 2B) on a range of low-dimensional dynamical systems that varied from 1-3 dimensions, which included multiple behaviours like chaos (Lorenz, Rossler, Sprott systems, etc. [27, 28]), multi-stability (pitchfork system), and oscillators (Van der Pol, Hopf normal form). In all cases, we trained the bias currents with gradient descent with momentum until a low loss was reached (Figure 2C). We found that the BANFF network could learn all the dynamical systems in the training set and immediately switch its output dynamics when a new learned bias set was applied (Figure 2D). This switching could occur across tasks and dimensions, with the last state of the network dictating the initial state of the network after a switch (Figure 2E, 2F). Note that, as the rank of the weights was static and fixed, when switching between a higher-dimensional task and a lower-dimensional task, superfluous dimensions could still be decoded out, but did not interfere with the target low-dimensional dynamical system. In some cases, we found that the initial condition upon a switch led to convergence to a spurious attractor (Supplementary Figure S1). We also note that for the chaotic attractors considered, the BANFF networks could readily reproduce either the chaotic attractor itself (Supplementary Figure S2) and the return map (Supplementary Figure S3), or heavily folded limit cycles that resemble the attracor near the edge of chaos (Supplementary Figure S2).

**Figure 2.**
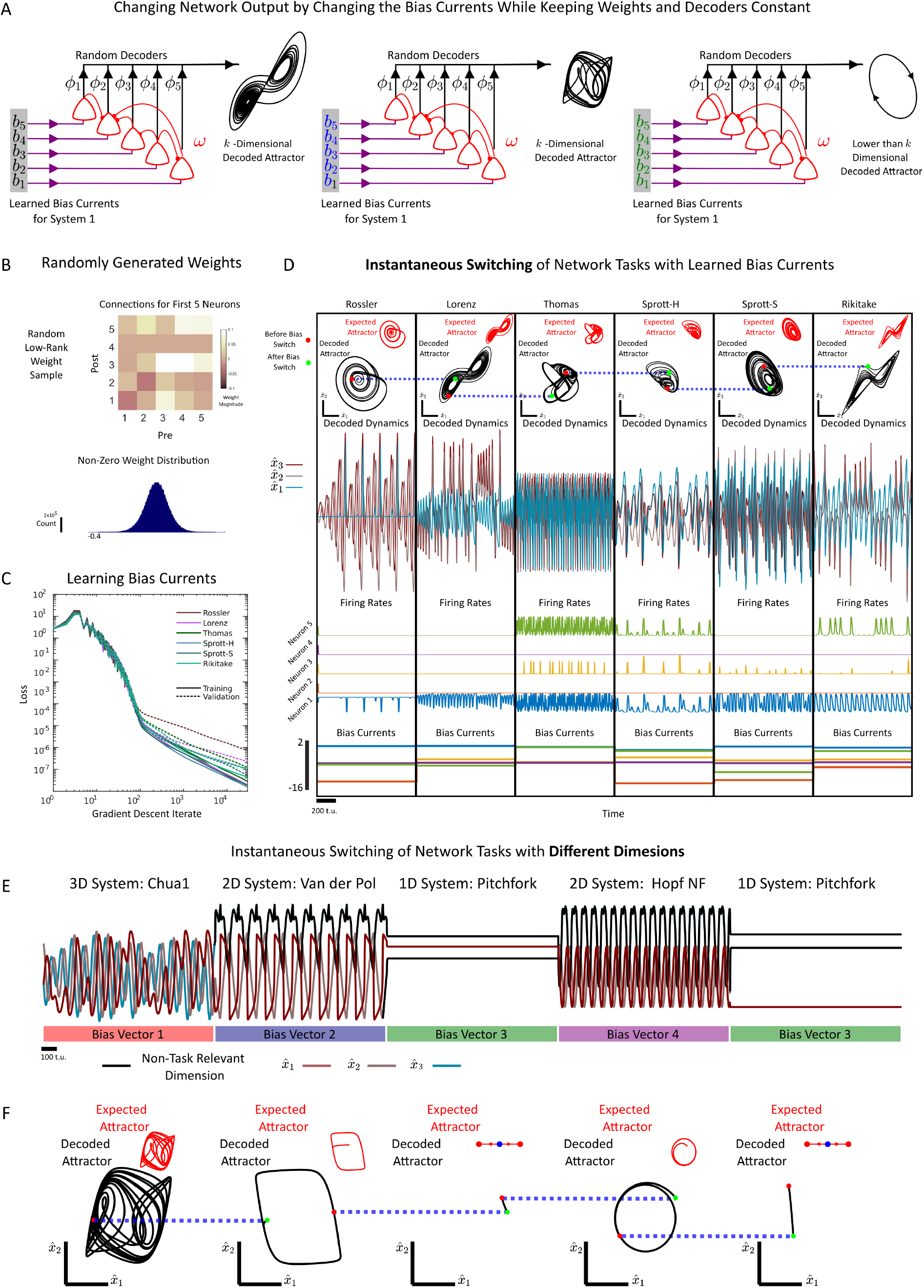
BANFF RNNs can Instantly Switch Between Different Dynamics by Learning Different Bias Currents. **(A)** A schematic for how a BANFF RNN with fixed weights can switch between multiple different dynamical systems. The learned bias currents (black, red, and blue) can be applied to the RNN to change the neuronal excitability and switch the task, in this case, switching between two different chaotic attractors and a lower-dimensional oscillatory system. **(B)** (Top) A selection of 5 fixed weights from the BANFF RNN. The weights for the low rank BANFF RNN are randomly generated and are approximately normally distributed (Bottom). **(C)** The loss versus the number of iterations of gradient descent with momentum for 6 separate chaotic systems. The training error (solid) and validation error (dashed) are shown with a total of 3 *×* 10^5^ iterations used during gradient descent. **(D)** The learned bias currents from (C) can be used to instantly switch between different tasks. (Top) The decoded attractors (in black) versus the expected attractors (in red). The red dots denote the last value of 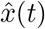 before the switch in the bias currents, while the green dots denote the value of 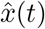 immediately after the switch in the bias currents. The decoded time-series for the dynamical systems under consideration are also shown (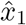, red, 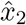, grey, 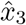, Cyan). The time-aligned firing rates for 5 neurons, and the time-aligned bias currents are shown in the bottom two rows. The bias currents are switched every 1000 time units (t.u.’s). **(E)** In a BANFF RNN, bias currents can switch between the dynamics of different dimensions. The learned bias currents are switched every 1000 t.u.’s to switch between 3D (Chua1 System), 2D (the Van der Pol and Hopf Normal Form systems) and 1D (the pitchfork normal form) dynamics. **(F)** The decoded (black) versus expected (red) attractor. The initial value of 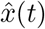 after a switch (green dot) and the final value before a switch (red dot) are plotted.

Finally, we investigated whether bias-only learning in BANFF networks was robust to biological constraints like Dale’s Law, where neurons forms exclusively excitatory or inhibitory connections to post-synaptic neurons (Supplementary Figure S4). To that end, we generated low-rank weight matrices that obeyed Dale’s Law by applying a simple constraint: all encoder components were constrained to be non-negative, and each neuron’s decoder components were required to share the same sign (Materials and Methods). As a result, every presynaptic neuron produced recurrent connections that were consistently either all excitatory (positive) or all inhibitory (negative). Next, we validated BANFF performance on 35 chaotic attractors (Supplementary Figure S4). Once again, a low-rank BANFF RNN with Dale’s law could mimic different dynamical systems, and instantly switch between them with the right bias current applied. We also note that despite the fact that the transfer function, tanh(*x*) has negative values, all learned low-rank BANFF networks and bias currents can be transformed into an equivalent network with a strictly positive firing rate (Supplementary Appendix 1).

In sum, low-rank BANFF RNNs were successfully validated on dozens of dynamical tasks without training any of the recurrent weights, even when the weights were randomly generated and biologically constrained. This implies that in order to learn multiple tasks, a recurrent neural network needs only to learn a series of bias currents to shift neural excitability in a task-specific way.

### BANFF Networks Provide a Mechanistic Explanation for Drifting Representations and Stable Computations with Noisy Units

We next investigated whether bias-only learning could account for representational drift, a commonly observed phenomenon in brain dynamics in which the activity patterns of individual neurons vary over trials or sessions while the overall task performance remains stable. Representational drift possibly reflects variability in hidden-layer activity without disrupting the functional dynamics required at the output level. [29,30, 31, 32, 33]. Bias currents adaptation provides a possible mechanism to account for representational drift, particularly if multiple bias currents implement identical task performance.

To test this, we serially trained a BANFF network on the Lorenz dynamical system with five separate bias current sets by initialising gradient descent with five separate initial conditions (Figure 3). We found that all five initial conditions led to successful learning, and that the bias current sets could be switched (Figure 3A) while the BANFF RNN was running. This produced no effect on the macroscopic behaviour of the system’s output (the Lorenz attractor), but immediately altered the activity patterns of individual neurons. On the long time scale, these bias currents could be interpreted as session-or day-specific. Neither the time series of the Lorenz system (Figure 3A) nor its phase portrait (Figure 3B) showed any indication of the bias-current switches, indicating that the output neurons continued to stably compute the correct macroscopic dynamics. Further, the bias currents were uncorrelated across neurons (Figure 3C), showing that excitability could change substantially after switching to a new bias set. In some cases, individual neurons were completely silent under one bias set (e.g., set ‘A’) yet strongly active under others (Figure 3A). This mirrored the results of *in vivo* long-time scale recordings, where neurons sometimes do and sometimes do not participate in a particular session of a task across multiple recording sessions [29, 30].

**Figure 3.**
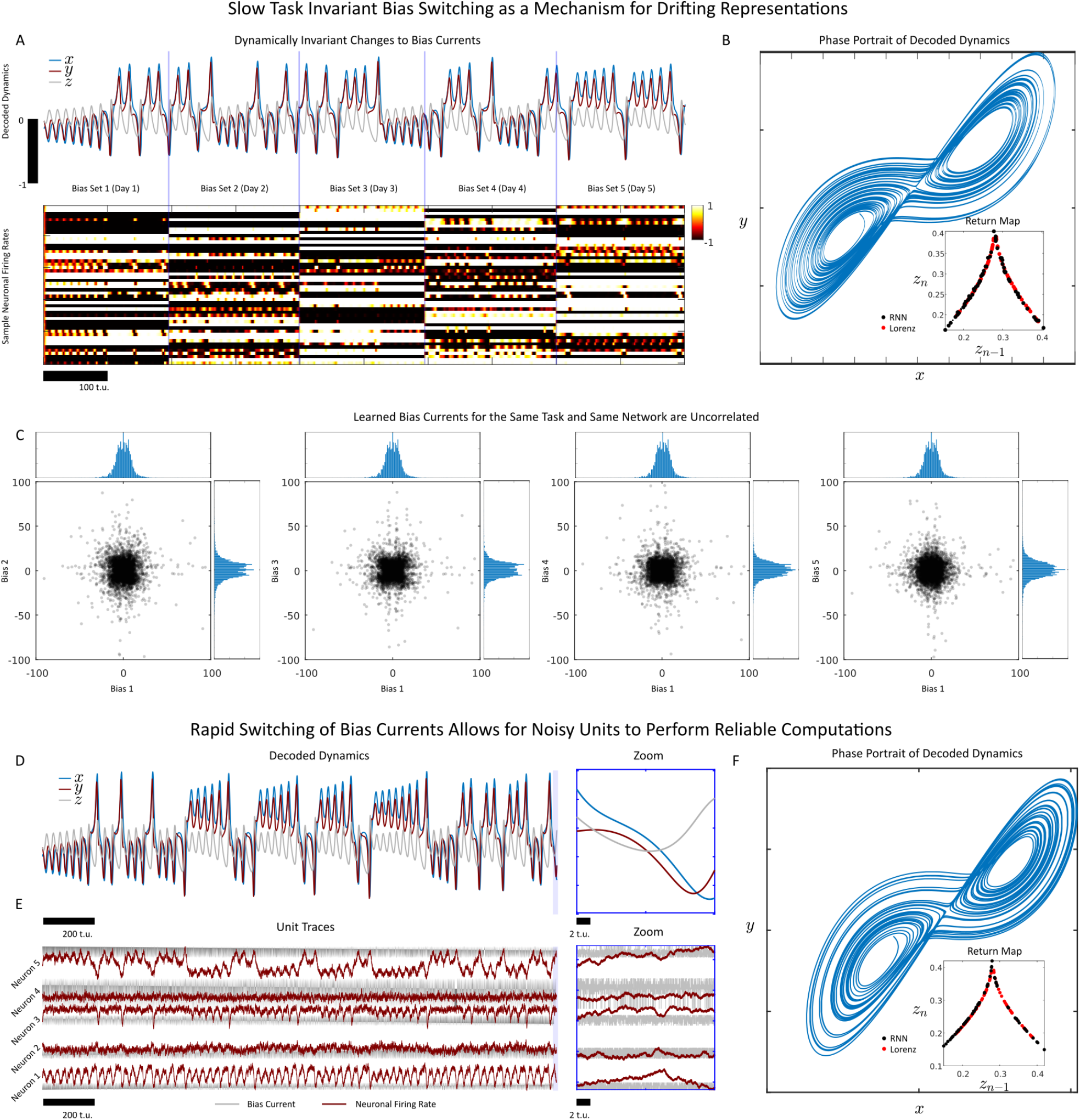
Non-Unique Learned Bias Currents for BANFF RNNs Provide Mechanisms for Representational Drift and Noise Robustness. **(A)** Multiple bias currents that allow a BANFF-RNN to mimic the same dynamical system are one potential mechanism to explain representational drift. Five separate bias currents were learned in order for a single BANFF RNN with fixed weights to mimic the Lorenz system. The 5 bias currents are switched every 200 time units, representing different days or recording sessions, as a model of drifting representation. **(B)** The phase portrait of the decoded Lorenz system. The return map is plotted as an inset for both the Lorenz system (red dots) and the BANFF RNN (black dots). **(C)** A plot of learned bias vector 1 correlated with bias vectors 2-5, along with the marginal distributions of bias vectors (on axes). The bias vectors display no obvious correlation and are uncorrelated. **(D)** The decoded dynamics for a BANFF RNN with noise-like bias currents. Despite the noisy dynamics of individual neurons (D), the decoded dynamics are substantially smoother. A zoom of the last 20 time units is shown (right). **(E)** The firing rates (red) for 5 RNNs with noise-like bias currents. The noisy inputs to these neurons were generated by switching between different learned bias currents at each time step. A zoom of the last 20 time units is shown (right). **(F)** The phase portrait of the decoded Lorenz system. The return map is plotted as an inset for both the Lorenz system (red dots) and the BANFF RNN (black dots).

We next examined how the network responded to rapid switches between different sets of bias currents (Figure 3D). Specifically, we considered the extreme case in which, at every time step, a single learned bias set was randomly selected and applied across all neurons. From the perspective of individual neurons, this input appeared as a highly variable, noise-like drive, even though it was generated by structured bias sets. Remarkably, this variability had no visible effect on the macroscopic dynamics of the system (Figure 3D), while individual neurons exhibited noisy activity as they filtered the rapidly changing bias currents (Figure 3E). The phase portrait also remained smooth (Figure 3F), showing no trace of the variability introduced by switching bias currents. Likewise, the Lorenz return map was unaffected. Moreover, in the supplementary materials (Supplementary Appendix 2), we provide a mathematical analysis that specifies the sufficient conditions under which neuron-level variability does not disrupt the macroscopic computation.

These results directly connect slow changes in neural representations (representational drift) to a possible source of noise-like variability during computation. Specifically, we predict that the correlated component of the noise a neuron receives on fast time scales drives the gradual reorganisation of representations over days. For example, if two neurons receive anti-correlated inputs on the millisecond-to-second scale, their resulting activity patterns mirror the divergence in activity that emerges across longer time scales during representational drift.

### Feedforward BANFF networks exhibit generalised learning capabilities

Although training bias currents to learn different dynamical systems demonstrates the generality of BANFF learning, these tasks primarily involve reproducing time series by learning relatively simple functions over state space that govern the system’s dynamics. We therefore asked whether bias-only learning could be applied more broadly to tasks relevant to both biological and artificial neural networks, and whether the same network could learn both dynamical and non-dynamical tasks—thus extending prior demonstrations of bias-only learning in static settings [17].

To test this hypothesis, we evaluated a feedforward BANFF network with two hidden layers, each comprising 16,000 neurons, across a set of 24 tasks spanning five categories: classification, regression, motor control, games and chaotic dynamical systems (Figure 4A-B). As in Figures 1-2, all weights were static and only the biases were changed using epoch-wise supervised training with the Adam optimiser [34]. For hidden layers, the outgoing weights adhered to Dale’s law to enhance the biological plausibility of the network. The number of input and output neurons in the network was task-specific, but the encoder/decoder weights were all sampled from identical weight matrices in the same order. All biases were initialised as zero, and a different set of biases was learned for each task (Figure 4C).

**Figure 4.**
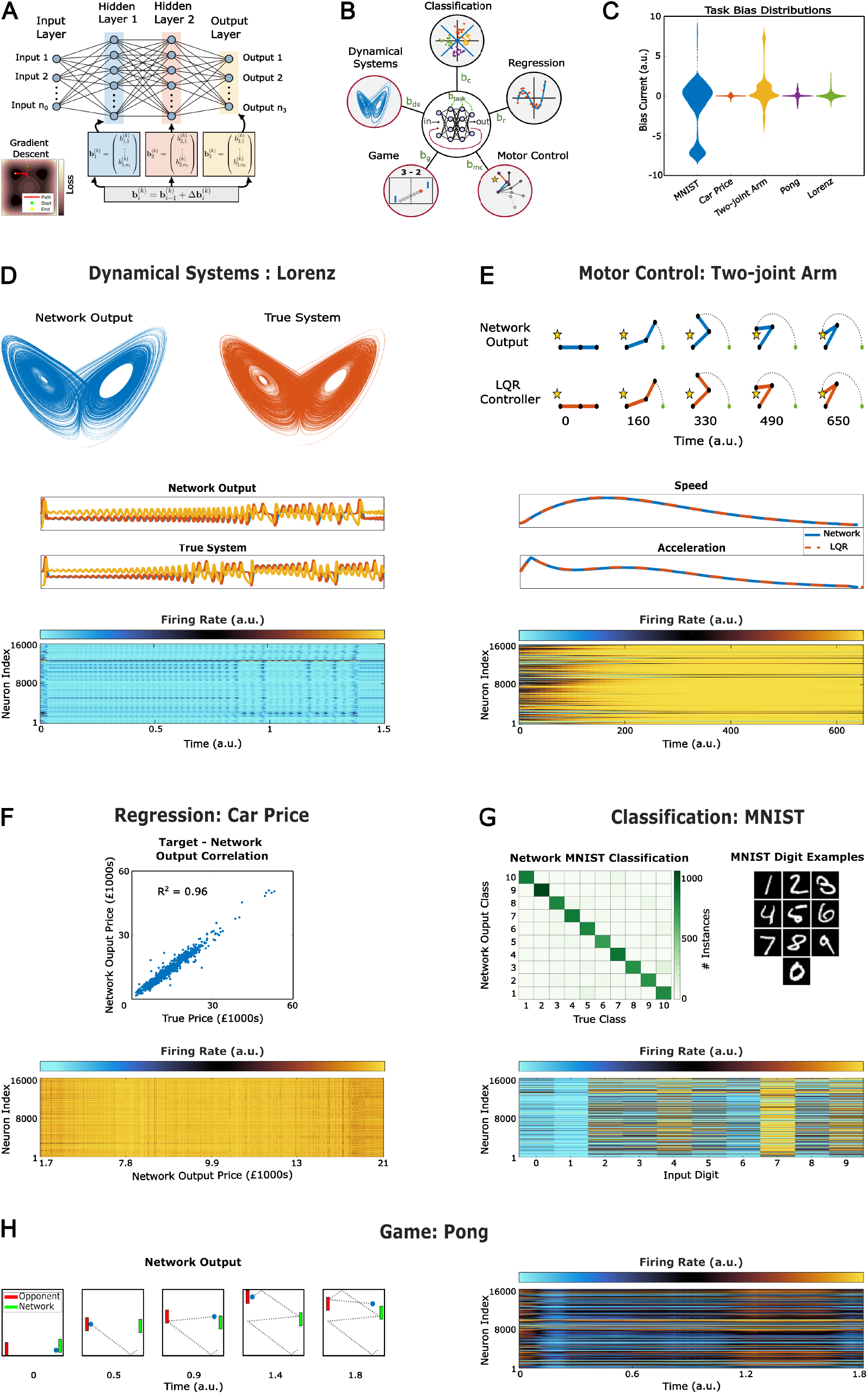
Generalised learning across tasks with a feedforward BANFF network. **(A)** A schematic of the structure of the feedforward network considered here. A gradient-based learning algorithm updates the bias inputs to the two hidden layers and the output layer. **(B)** The five classes of tasks learned by the network. Black and red outlines indicate that the network was tested in open- and closed-loop configurations, respectively. **(C)** The neuronal bias distributions for the second hidden layer of the network (16,000 neurons) for one example task belonging to each class of task. **(D)** (Top) The phase portraits for the output of the network when trained to emulate the Lorenz system in a closed-loop mode alongside the true phase portrait visualised using the deterministically simulated system. (Middle) The time series output from the network and the true system, each colour represents one dimension of the three-dimensional output. (Bottom) The activity of all neurons in the second hidden layer of the network over the same time frame as the output shown above. **(E)** (Top) An example trajectory of the two-joint arm performing a reaching task when controlled by the network and the LQR controller, respectively. (Middle) The speed and acceleration of the end point of the two-joint arm over the same example trajectory. (Bottom) The firing rate of all neurons in the second hidden layer over the same example trajectory. **(F)** (Top) A correlation between the true price and network output price for a car price regression task. (Bottom) The firing rate of all neurons in the second hidden layer of the network as a function of the network output price. **(G)** (Top left) A confusion matrix for the network output and true class for the MNIST task. (Top right) Example images of each digit for the MNIST task. (Bottom) The firing rate for all neurons in the second hidden layer of the network for the input images shown in the top right of the panel. **(H)** (Left) An example trajectory of pong play showing the network controlling one paddle and an optimal player controlling the other. (Right) The firing rates of all neurons in the second hidden layer of the network across the same time frame as the trajectory shown to the left.

For motor control, the game Pong, and chaotic dynamical systems tasks, the network was trained in an open-loop mode using custom-generated synthetic training datasets. The network was then tested in a closed-loop configuration, whereby its output was recursively fed back as input. This configuration allowed the output errors to accumulate as they were recursively processed by the network, effectively ensuring that the network had to emulate the target dynamics successfully and not just ‘memorise’ the supervisor to perform well.

Similarly to the RNN, the feedforward BANFF network was employed to learn the dynamics of nine chaotic dynamical systems tasks. In each instance, the network was trained in an open-loop mode to predict the next state of the system given the current state as an input. The target for learning was a deterministically simulated time series of the system. For the subsequent closed-loop testing, simulation began with a unique initial state not used in training, and no information about target dynamics was provided. The network reliably reproduced characteristic phase portraits and return maps that were visually congruent with those of the underlying deterministic systems (Table 1, Figures 4D, Supplementary Figures 5 & 6 and Supplementary Video 1). The output time series of the network was not identical to the deterministically simulated system, which was expected because the target systems were chaotic. More importantly, the network was able to replicate the phase portraits and return maps of the systems, demonstrating that the intrinsic dynamics of the system had been learned [35].

**Table 1.**
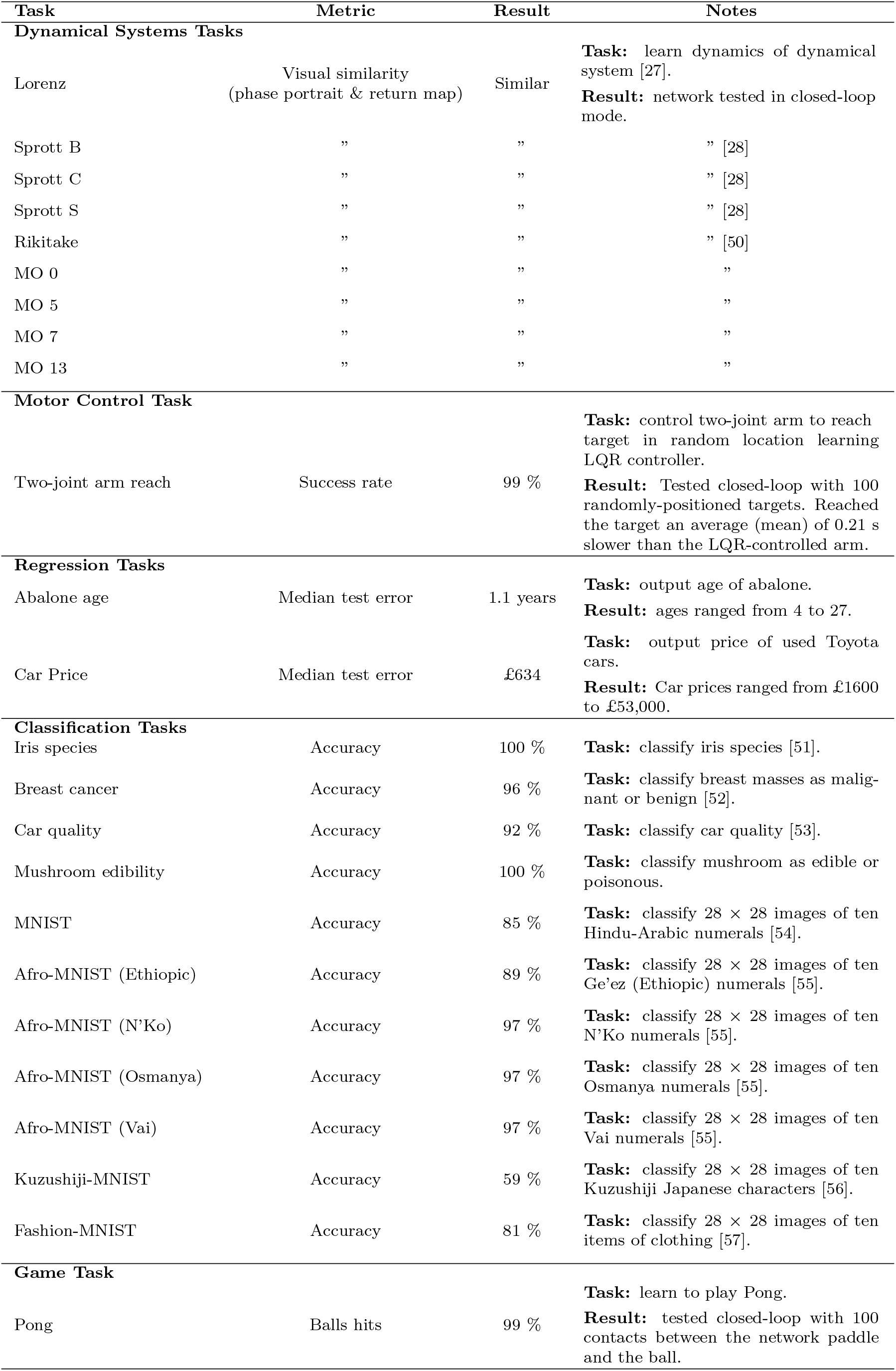
The numerical performance of the feedforward BANFF network across all tasks. Further details may be viewed in the supplementary material.

| Task | Metric | Result | Notes |
| --- | --- | --- | --- |
| **Dynamical Systems Tasks** |  |  |  |
| Lorenz | Visual similarity<br>(phase portrait & return map) | Similar | **Task:** learn dynamics of dynamical system [27].<br>**Result:** network tested in closed-loop mode. |
| Sprott B | ” | ” | ” [28] |
| Sprott C | ” | ” | ” [28] |
| Sprott S | ” | ” | ” [28] |
| Rikitake | ” | ” | ” [50] |
| MO 0 | ” | ” | ” |
| MO 5 | ” | ” | ” |
| MO 7 | ” | ” | ” |
| MO 13 | ” | ” | ” |
| **Motor Control Task** |  |  |  |
| Two-joint arm reach | Success rate | 99 % | **Task:** control two-joint arm to reach target in random location learning LQR controller.<br>**Result:** Tested closed-loop with 100 randomly-positioned targets. Reached the target an average (mean) of 0.21 s slower than the LQR-controlled arm. |
| **Regression Tasks** |  |  |  |
| Abalone age | Median test error | 1.1 years | **Task:** output age of abalone.<br>**Result:** ages ranged from 4 to 27. |
| Car Price | Median test error | £634 | **Task:** output price of used Toyota cars.<br>**Result:** Car prices ranged from £1600 to £53,000. |
| **Classification Tasks** |  |  |  |
| Iris species | Accuracy | 100 % | **Task:** classify iris species [51]. |
| Breast cancer | Accuracy | 96 % | **Task:** classify breast masses as malignant or benign [52]. |
| Car quality | Accuracy | 92 % | **Task:** classify car quality [53]. |
| Mushroom edibility | Accuracy | 100 % | **Task:** classify mushroom as edible or poisonous. |
| MNIST | Accuracy | 85 % | **Task:** classify $28 \times 28$ images of ten Hindu-Arabic numerals [54]. |
| Afro-MNIST (Ethiopic) | Accuracy | 89 % | **Task:** classify $28 \times 28$ images of ten Ge'ez (Ethiopic) numerals [55]. |
| Afro-MNIST (N'Ko) | Accuracy | 97 % | **Task:** classify $28 \times 28$ images of ten N'Ko numerals [55]. |
| Afro-MNIST (Osmanya) | Accuracy | 97 % | **Task:** classify $28 \times 28$ images of ten Osmanya numerals [55]. |
| Afro-MNIST (Vai) | Accuracy | 97 % | **Task:** classify $28 \times 28$ images of ten Vai numerals [55]. |
| Kuzushiji-MNIST | Accuracy | 59 % | **Task:** classify $28 \times 28$ images of ten Kuzushiji Japanese characters [56]. |
| Fashion-MNIST | Accuracy | 81 % | **Task:** classify $28 \times 28$ images of ten items of clothing [57]. |
| **Game Task** |  |  |  |
| Pong | Balls hits | 99 % | **Task:** learn to play Pong.<br>**Result:** tested closed-loop with 100 contacts between the network paddle and the ball. |

In the motor control task, the network controlled a two-joint arm to reach a randomly positioned target (tested using 100 targets), with supervisory signals provided by a linear quadratic regulator (LQR) controller. During closed-loop testing, the state of the arm was iteratively updated based on the network’s output, and the system achieved target acquisition in 95% of trials, demonstrating successful emulation of the LQR controller (Table 1, Figure 4E and Supplementary Video 1).

For classification and regression tasks, the network was trained and evaluated in an open-loop configuration using previously unseen test datasets. Under these conditions, the network successfully learned all tasks, demonstrating a test performance comparable to those reported by Williams *et al*. (Table 1 and Figure 4F & G) [17]. The slightly lower performance was expected, given that enforcing biological plausibility imposed constraints on the network at the cost of task performance.

In a dynamic game setting, the network was tasked to control a paddle in the classic arcade game Pong. During training, the network was supervised to re-centre its paddle after ball contact and subsequently to align its vertical position with the approaching ball after the opponent’s return, once the ball was sufficiently close. During closed-loop testing, wherein the paddle and ball’s position were continuously fed back as input, the network demonstrated successful emulation of the supervisor, attaining a 99 % hit rate when simulated against an error-free opponent (Table 1, Figure 4H and Supplementary Video 1).

The bias distributions for each task were distinct and varied in their respective deviation from the initial distribution (Figure 4C). This was also clear from the observed activity of the neurons, which showed variability between tasks (Figure 4D-H). For example, the Lorenz dynamical system task resulted in an overall shift towards negative bias, lowering the firing rate of most neurons, whereas the regression task resulted in biases that closely matched the initial distribution. As a result, the firing rates of neurons for this task were generally larger.

Collectively, these results demonstrate the efficacy and versatility of tuning neuronal excitability as the basis for learning. This was the case for both RNNs and feedforward networks operating in both open-and closed-loop task configurations, providing evidence that such architectures may present a candidate functional model of some aspects of brain computation via control of neural dynamics.

## Discussion

Since the time of Donald Hebb, it has been posited that the main form of learning in the brain arises from changes in the synaptic connections between neurons [36]. In this study, we have demonstrated that recurrent and feedforward networks with fixed and random synaptic connectivity can perform multiple complex tasks by learning a task-specific bias current for each neuron. In this learning paradigm, which we term the Bias Adaptive Neural Firing Framework (BANFF), networks perform a wide range of tasks, from image classification to dynamical system forecasting and motor control, providing a parsimonious and biologically plausible account of one potential mechanism of learning in the brain. This observation is consistent with recent studies suggesting that learning in the brain can occur without synaptic plasticity, but rather with learned upstream inputs or plasticity in the intrinsic excitability of neurons themselves [37]. Learning in BANFF networks is efficient both because of the reduced number of trainable parameters and because new tasks can be acquired without interfering with previously learned ones. Moreover, BANFF networks readily operate under important biological constraints (e.g., Dale’s Law), and the learning rule remains largely local when the recurrent weight matrix is low-rank.

The theoretical framework developed by Williams *et al*. [17] shows that neural networks retain universal approximation capabilities even when neuron biases are trained and synaptic weights remain fixed and random. Related work in reservoir computing has demonstrated that fixed inputs with adaptable weights can similarly drive transitions between distinct low-dimensional dynamics [38]. In addition, previous studies have linked changes in neural excitability to both computation and representational drift [39, 37, 40].

Our findings extend these principles, demonstrating bias-only learning and its applicability beyond open-loop feedforward function approximation, encompassing the dynamic, closed-loop temporal processing necessary for motor control, game play, and dynamical systems modelling. Thus, bias modulation alone can effectively reconfigure network dynamics in behaviourally meaningful ways. We also show that switching bias currents on long time scales can account for representational drift, while switching them on short time scales provides a mechanism for robust computations with noisy units. Critically, BANFF networks predict that noise-robust computations are mediated by the same mechanism as representational drift: changes to neuronal excitability, with the only mechanistic difference between the two being the time-scale at which bias currents switch, which is also in line with recent previous work that implicates excitability changes as the mechanism behind representational drift in spiking network models [40].

One compelling biological interpretation of the adjustable biases in our BANFF networks is that they represent external inputs from subcortical structures, such as the basal ganglia, whose role in cortical modulation is extensively documented [7, 8]. This interpretation aligns with the notion of *outsourced* cortical learning, wherein subcortical structures adjust cortical responses through neuromodulatory signals or persistent input patterns [7, 8]. Such a mechanism has been well-explored in basal ganglia-cortical models of motor learning, where basal ganglia outputs influence cortical activity patterns to facilitate rapid behavioural adjustments and stabilisation. Crucially, this external bias modulation could address the inherent instability issues associated with other plasticity mechanisms, which rely on synaptic changes (i.e., spike-timing-dependent plasticity), that can lead to instability in recurrent networks [41]. Bias-based modulation via subcortical signals may be able to rapidly stabilise and finely tune cortical network dynamics without extensive and potentially disruptive synaptic remodelling [41]. This outsourced model, therefore, provides a biologically plausible solution to the challenge of achieving rapid, stable learning in neural circuits without compromising the underlying network integrity.

Furthermore, evidence suggests that intrinsic neuronal excitability serves as another non-synaptic mechanism for modulating cortical processing in the short term. Intrinsic excitability changes, involving modifications in spike thresholds and resting membrane potentials, have been documented *in vivo* as mechanisms for adapting neural responses without altering synaptic strength [42, 43]. Additionally, BANFF networks inherently mitigate the risk of catastrophic forgetting, which often plagues networks trained sequentially on multiple tasks [44] as synaptic connectivty can remain fixed while tasks change. Further, the bias currents trained by BANFF learning can be stored in simple network structures (e.g. winner-take-all networks) where the number of stored tasks scales linearly with the network size. This property aligns closely with neuroscientific observations, suggesting that neuromodulatory and excitability-based mechanisms allow the brain to rapidly reconfigure its functional circuits to meet diverse, context-specific demands without significant structural changes.

In summary, our work demonstrates that learning through bias modulation alone represents a robust and biologically plausible mechanism for rapid behavioural adaptation that is not exclusive of other processes, such as spike time-dependent plasticity. The bias-learning paradigm aligns with experimental evidence suggesting the importance of bias-like mechanisms in cortical processing, and it advances theoretical perspectives on neural network expressivity.

## Acknowledgements (not compulsory)

WN is funded by an NSERC Discovery Grant (RGPIN/04568-2020), a Canada Research Chair (CRC-2019-00416), a Hotchkiss Brain Institute start-up grant and the Cumming Medical Research Fund. AG is supported by NSERC, Digital Research Alliance of Canada, Hotchkiss Brain Institute, Alberta Children’s Hospital Research Institute, and the Azrieli Accelerator. This research was partially funded by Synaptrain Technologies Inc.

## Author contributions statement

WN and AG supervised the project. WN, KM, performed numerical simulations. WN, KM performed mathematical analysis. WN, AG, CC, KM, and SS prepared and edited the manuscript.

## Code Availability Statement

The code for this project will be made available at the following link: https://github.com/kaimason100/BANFF_ANN

## Methods

### Recurrent BANFF Networks

The recurrent neural networks (RNNs) considered here had the general form:

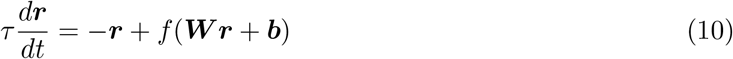

where ***r***(*t*) was an *N ×* 1 dynamical vector of filtered neuronal activity, and ***W*** was the *N × N* matrix of synaptic weights. The function *f* (*z*) was assumed to be a saturating neuronal transfer function. In this work, we exclusively used

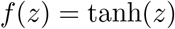

The *N ×* 1 vector ***b*** represents neuronal bias currents, used as free parameters for learning target dynamics. Further, a set of readout weights, ***ϕ***, where ***ϕ*** is a *N × k* matrix, was used to decode the dynamics 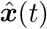 using the filtered rates ***r***:

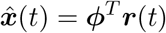

In training a typical RNN, the readout weights ***ϕ*** and the recurrent weights ***W*** are used as free parameters to minimise some loss or cost function, 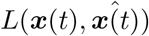, that compares ***x***(*t*), the dynamics of a target system, with 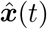, the decoded output of the RNN. For example, in backpropagation-through-time, all weights (***W***, ***ϕ***) and bias currents ***b*** are used as learning parameters, with the gradients 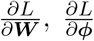, and 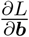 computed to iteratively update the weights and biases.

A BANFF network simplifies both the learning rules and network structure from the general RNN equations (10)-(11) in two ways. First, we considered low rank weights. In particular, ***W*** was assumed to be of rank *k*, where *k* was small and decomposed as the following outer product

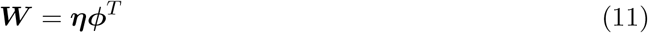

This is a similar weight structure to the Neural Engineering Framework [21, 22, 26] with the key difference being that in the NEF, bias currents are generated randomly, and decoders (***p****hi*) are learned, rather than the approach here, which is to learn bias currents and generate decoders randomly. The matrix ***η*** was *N × k* and served to encode the target dynamics 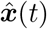 as:

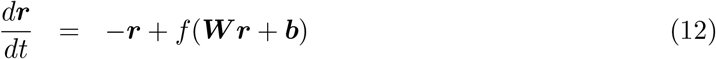

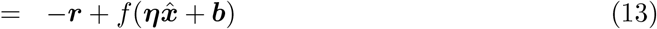

Further, with the assumption (11), then the dynamics of 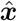 were given by

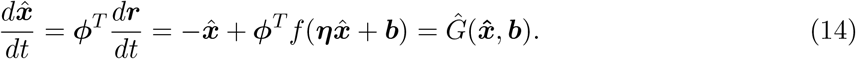

The ***η*** and readout weights, ***ϕ***, were randomly generated, but from constrained distributions that conferred functionality and fast learning to the networks. The first constraint was on the components of ***η***, *η*_*ij*_ for *i* = 1, 2, … *N* and *j* = 1, 2, … *k* were drawn from a ternary distribution consisting of *{−M*, 0, *M}*. This ternary distribution, whilst not necessary when the network was constrained to *k*-dimensional dynamics, was crucial for allowing the network to flexibly represent dynamical systems that were less than *k*-dimensional, even if the rank of ***W*** is *k*. In short, this ternary distribution in the components of ***η***_*ij*_ ensured that a random subset of neurons would exclusively encode at most *l < k* variables of 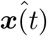.

Next, the readout weights ***ϕ*** were constrained in how they scaled with the network size:

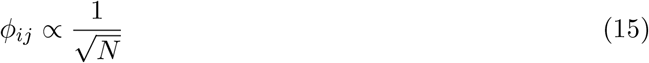

As this led to faster learning, despite the fact that the encoders and decoders were held fixed.

### Learning in recurrent BANFF networks

With the constraints on the rank of ***W***, equation (11), in addition to the constraints on ***η*** and ***ϕ***, the network outputted a dynamical system with the following form:

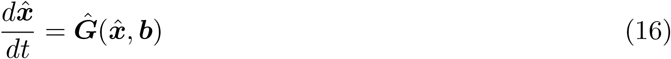

where ***b*** was the vector of *N ×* 1 bias currents that parameterised the function *F* .

If the dynamics of ***x***(*t*) (the supervisor) satisfy the general non-linear non-autonomous dy-namical system:

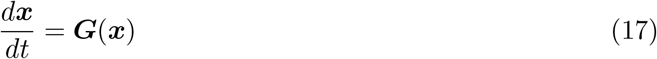

Then one can train the dynamics of 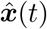 to mimic the dynamics of ***x***(*t*) by using the bias vector ***b*** to minimise the following loss function, *L*(***b***)

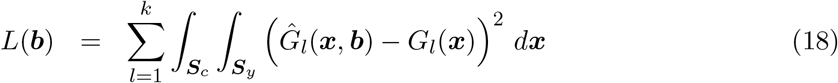

where

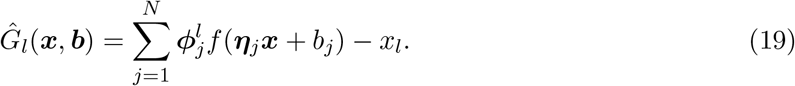

This loss function was, in practice, always replaced with the discrete loss:

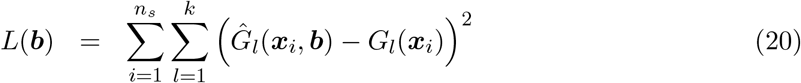

where *n*_*s*_ was the number of times the dynamics of the supervisor had been sampled.

To learn a vector of bias currents ***b*** that minimised equation (18), we first computed the gradient with respect to the bias vector:

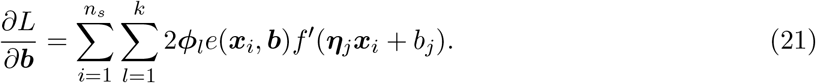

With the gradient of the loss with respect to the bias currents computed, any gradient descent-based learning rule could be used to update ***b***. A basic gradient descent with momentum rule was used to learn ***b***

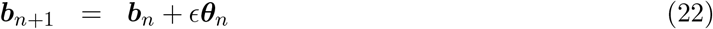

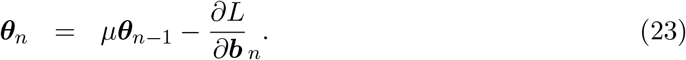

### Dynamical Supervisors Used to Train the BANFF RNN

All systems used to train the BANFF RNNs were *k*-dimensional autonomous dynamical systems of the form

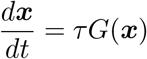

where *τ* controlled the time-scale of the supervisor. To train the BANFF RNNs, each system was simulated for 1000 time units. This simulation sampled the attractor of the system so that the system could also be rescaled and centred in space with:

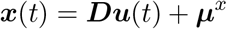

where

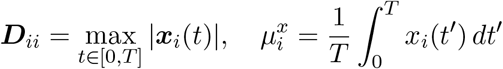

The BANFF RNN was then trained to approximate the function. A series of points, ***u***_*i*_, *i* = 1, 2, … *n*_*u*_ were then randomly drawn from the *k* dimensional cube [*−*1, 1]^*k*^. The BANFF RNN was then trained to mimic the dynamics of 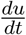 by learning the function:

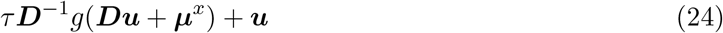

The dynamical systems considered can be found in Supplementary Tables 1-5, along with the initial conditions used to simulate ***x***(*t*) for each supervisor, and estimate ***D*** and ***µ***^*x*^. By learning Equation 24 through modifying bias currents, BANFF RNNs mimic

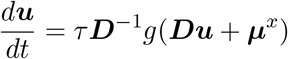

which is the rescaled version of 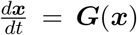. Note that all figures utilize ***u***(*t*), the rescaled dynamics.

### Generalised Learning with Feedforward BANFF Networks

#### Network Structure

We used one fully connected feedforward NN comprising an input layer, two hidden layers, and an output layer (Figure 4). We varied the number of neurons in the input and output layers to suit the task at hand, and the hidden layers always comprised 16,000 neurons each. The weights between neurons were fixed for all tasks and were initialised such that the outgoing weights of neuron *j* in the *i*^th^ layer

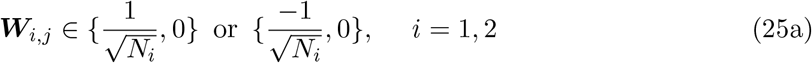

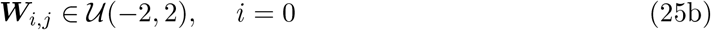

where *N*_*i−*1_ denotes the number of neurons in the (*i −* 1)^th^ layer (*i* = 0 corresponds to the input layer) [45, 46]. Neurons with output weights corresponding to (25a) were dubbed excitatory and inhibitory, respectively [46]. Whether a neuron was excitatory or inhibitory was random with probability 1/2. Whether an outgoing weight of a neuron was non-zero was random with probability 2/3, such that the distribution of weights across all neurons was uniform. ***W***_0_, the encoding weight matrix, did not conform to this distribution, but was sampled (in the same order for each task) from a single encoding matrix which was itself sampled from a random uniform distribution between -2 and 2 (Equation 25b). This was chosen since it was an encoding layer from the inputs to the first layer of hidden neurons, and the uniform distribution created a sufficiently rich encoding. For all tasks, the inputs to the network were scaled by 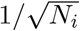 where *N*_*i*_ was the number of inputs.

The activation (output) of the *i*^th^ layer of the NN was

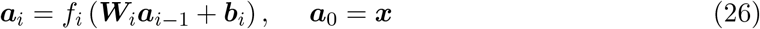

where *f*_*i*_(*z*) was the activation function of the *i*^th^ layer, ***W***_*i*_ was the weight matrix with size *N*_*i*_ *×N*_*i−*1_, ***a***_*i−*1_ was the activation with size *N*_*i−*1_ *×* 1, ***b***_*i*_ was the bias with size *N*_*i*_ *×* 1 and ***x*** was the input with size *N*_1_ *×* 1. The activation functions were

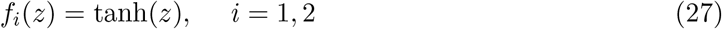

and

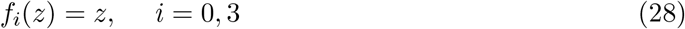

for non-classification or game tasks, or

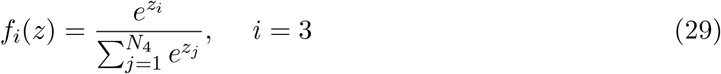

(the softmax function) for classification and game tasks [47].

#### Network Training

We trained the network in a supervised fashion using the Adam optimiser (epoch-wise) with backpropagation to change the biases for all tasks. All weights remained static [48, 34, 17]. The update rule for the bias of neuron *j* in the *i*^th^ layer of the network for epoch *k* was

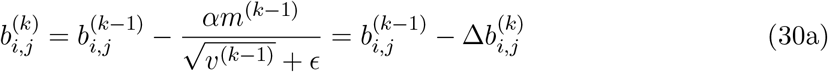

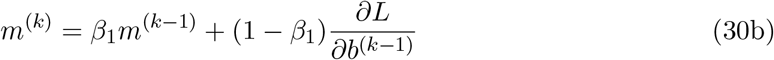

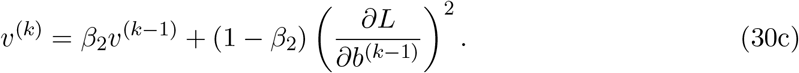

*α* was the learning rate and *β*_1_, *β*_2_ and *ϵ* were hyperparameters heuristically chosen as 0.9 0.999 and 1 *×* 10^*−*8^ for non-dynamical systems tasks and 0.99, 0.999 and 5 *×* 10^*−*6^ for dynamical systems tasks, respectively [34]. Here, *β*_1_ and *β*_2_ were exponential decay rates that control how past gradient information contributed to the first- and second-moment estimates, respectively, with larger values giving smoother but slower adaptation. The constant *ϵ* was a small positive term added for numerical stability to prevent division by zero [34]. *L* was the loss function, which was defined as either the mean-squared-error loss *L*_*MSE*_ or the cross-entropy loss *L*_*CE*_. The mean-squared-error loss was used for all non-classification/game tasks and was defined as

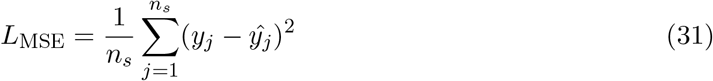

where *n*_*s*_ was the number of samples in the epoch, *y*_*j*_ was the *j*^th^ element of the supervisor and 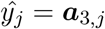 was the output of the network for the *j*^th^ output neuron. The cross-entropy loss was used for all classification and game tasks and was defined as

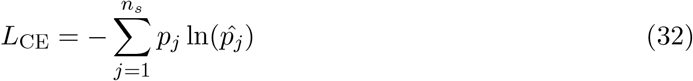

where *p*_*j*_ was the *j*^th^ element of the supervisor and 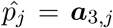 was the *j*^th^ element of the network output.

#### Tasks

A total of 24 tasks belonging to five categories were learned (Figure 4). For each task, the data were split into training, validation and test datasets.

For all classification tasks, all data were taken from pre-existing datasets, and the inputs were features pertaining to the task. For MNIST tasks, the inputs were 28*×*28 images, and each feature was a pixel intensity. For other classification tasks, the features were numerical measurements or categorical classes. The network’s performance was assessed based on the percentage of correctly classified test cases (% accuracy).

For the motor control task, the network was trained to imitate a linear–quadratic regulator (LQR) that controls a planar two-link arm in a 2-D reaching task. The arm had link lengths of 1.0 m and 0.8 m with joint angles *θ*_1_ and *θ*_2_. The state vector was

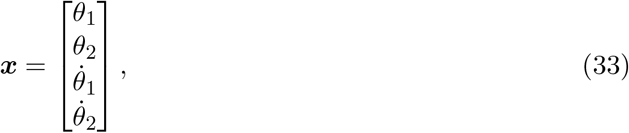

and the dynamics were modelled as

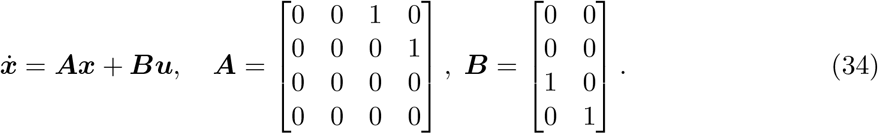

The LQR minimized

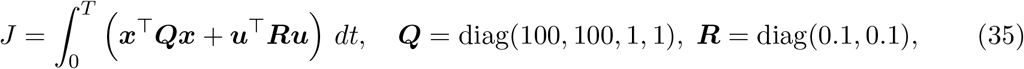

with optimal feedback ***u*** = *−****K***(***x*** *−* ***x***^*∗*^), where 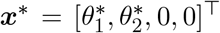 is the desired joint state obtained by inverse kinematics for the sampled end-effector (EE) target, and ***K*** is given by the continuous-time algebraic Riccati equation solution.

Each episode started from ***x***_0_ = **0**. A random reachable EE target 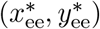 was sampled, converted once to the desired joint state ***x***^*∗*^ (elbow-up branch, with the usual clipping of cos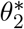 to [*−*1, 1]). The LQR was rolled out for up to *T* = 5 s with Δ*t* = 0.01*s*, and the rollout terminated early for data collection when the EE distance fell below 0.01 m for at least 10 consecutive steps (0.1 s). At each step, the input to the network was a six-dimensional *joint-error* feature vector

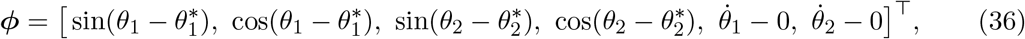

and the target output was the corresponding LQR control ***u***. We generated 250 such episodes; an episode-level split (80%/20%) was used for training/validation to avoid trajectory leakage.

Testing used 100 novel targets and identical initial conditions for the arm. The controller operated in closed loop: at each time step, the current state ***x***_*k*_ produced the joint-space error ***e***_*k*_ = ***x***_*k*_ *−* ***x***^*∗*^, which was encoded as ***ϕ***_*k*_, passed through the network to produce ***u***_*k*_, and applied to the plant to obtain ***x***_*k*+1_. The target joint state ***x***^*∗*^ remained constant for the episode. Success required the EE to be within 0.1 m of the target for at least 10 consecutive steps (0.1 s) within *T* = 5 s. For fair comparison, both the LQR and the network were simulated on the same time grid, with terminal-state padding to equalise horizon length.

For the Pong task, we generated a training and validation dataset by randomly sampling ball and paddle positions along with the ball’s velocity. These six quantities (the *x*- and *y*-components of each) were provided as inputs to the network. Training was supervised: the network was taught first to return its paddle to the centre after hitting the ball, and then to track the ball’s *y*-position once the opponent had returned it and it crossed a defined threshold of closeness (*x* ≥ 0.7 in a game domain with *x ∈* [0, 1]).

The problem was framed as a classification task in which, at each time step, the network produced a 1 *×* 3 one-hot output vector indicating whether the paddle should move up, move down (each by a fixed discrete distance), or remain stationary. For testing, the trained network played against an opponent whose paddle always matched the ball’s *y*-position. Performance was quantified by recording the number of missed balls across 100 paddle–ball contacts.

For regression tasks, pre-existing datasets were used, and each dataset was split into training, validation and test data. The inputs corresponded to features pertaining to the system under investigation, and the outputs corresponded to predicted quantities. The network’s performance was assessed on the median error between its predictions and the true value for the test dataset in each case. The error was defined as |*y−ŷ*| where *y* was the true value and *ŷ* was the network’s output.

For dynamical systems tasks, the dynamical system being learned was simulated deterministically using the `ode45` function in MATLAB up to a time of 3,000 time units. This was used as a training/validation dataset by using the *n*^th^ values of the time series as the supervisor and the (*n −* 1)^th^ values of the time series as inputs. For testing, the network was given an initial condition equal to that of the training data plus a random perturbation sampled from [0, 0.02] and then simulated in a closed-loop configuration (i.e. the network’s output was used as the value for the *n*^th^ time step for future predictions) for a further 20,000 time units. The network’s output was compared to the true system by visual comparison of the phase portraits and return maps.

#### Implementation

All simulations were performed on a desktop computer (**CPU**: 3.80 GHz AMD Ryzen 7 5800X (8-Core); **RAM:** 64 GB; **GPU:** NVIDIA GeForce RTX 3050) using Matlab and the Deep Learning Toolbox for neural network implementations [49]. The computer’s GPU was utilised for neural network training.

## Additional Methods

### Additional Methods for Figure 1

For Figure 1E, a network of 50 uncoupled tanh neurons was used to approximate the function *y* = *x*^2^, using gradient descent on the bias currents with a learning rate of *ϵ* = 0.1. The decoders were randomly generated from a normal distribution with mean 0, and standard deviation 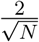. Identical parameters were used for Figure 1F with the supervisor *y* = cos(2*πx/*10). For Figure 1F, the encoder components were randomly generated and sparse, with:

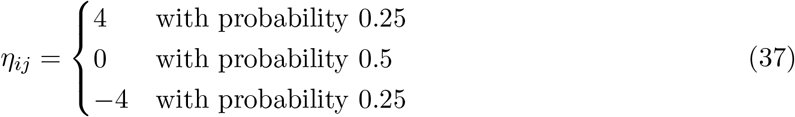

while the encoders were drawn from a normal distribution with mean 0, and standard deviation 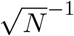 with *N* = 6000. Gradient descent with momentum was used to learn the bias currents, with the learning rate set to *ϵ* = 0.1 and the momentum parameter set to *γ* = 0.9. In Figure 1J, the learned bias currents from Figure 1I were “turned on” at time *t* = 1000 t.u. (arbitrary time units), with the bias currents all set to 0 for *t <* 1000 t.u. The RNN was integrated with a Runge-Kutta integrator in MATLAB (`ode45`).

### Additional Methods for Figure 2

All networks considered in Figure 2B-E, contain *N* = 6000 neurons with randomly generated encoders and decoders as in Figure 1I-K. Owing to the low-rank RNN structure, the resulting network weights were determined by ***W*** = ***ηϕ***^*T*^ . The distribution of non-zero weights is plotted in Figure 2B. To learn the bias currents, gradient descent with momentum was used with identical parameters, as shown in Figure 1I-K. The systems in Figure 2E-F were integrated for *t* = 300 t.u.’s with ode45.

### Additional Methods for Figure 3

The same BANFF RNN with *N* = 6000 neurons (as in Figure 1-2) was trained to mimic the Lorenz system with 20 different bias currents. The bias currents were all initialised with different initial conditions to force the network to find different solutions. The bias currents were initialised with a random normal distribution with a mean of 0 and a standard deviation of 3. Each bias current underwent gradient descent with momentum with a momentum parameter of *γ* = 0.9, and an initial learning rate of *ϵ* = 0.1 for 10^4^ iterations. Figure 3A-B were simulated with `ode45`, for a total of 2000 time units, with the bias current being switched every 400 time units between the first five learned bias currents. In Figure 3D-F, the network was integrated with a basic forward Euler integration scheme with an integration time constant of Δ = 10^*−*2^ time units. At every time step, the bias current was randomly selected from one of the 20 learned bias currents. Euler integration was used in D-F as the simulation time was much quicker with rapidly switching bias currents.

## Additional Methods for Figure 4

The network considered for all tasks in Figure 4 comprised two hidden layers of 16,000 neurons each. For Figure 4 **a**, a generic gradient descent path is shown, not corresponding to any task in particular. For Figure 4 **c**, only the biases of the second hidden layer are shown for a subset of the tasks considered. For Figure 4 **d**, the Lorenz system was initialised with a small random perturbation from the initial state used in training. The true system and the network were both simulated for 7,000 steps with the network working in closed-loop mode. For Figure 4 **e**, the target was placed in a random location and the network was simulated in closed-loop mode to control the two-joint arm to reach the target (the position of the target was still fed open-loop). For Figure 4 **f**, the network was tested on a hidden test set of data and the network’s output was compared to the true values. For Figure 4 **g**, the network was tested on a hidden set of test data and the classification results plotted. The firing rates shown correspond to the numbers shown in the top right of the panel. For Figure 4 **h**, the network was simulated in closed-loop mode to play pong against a perfect opponent. In all panels, the firing rates shown are those of the second hidden layer of neurons in the network.

## Additional Methods for Supplementary Figure S1

Each BANFF RNN and supervisor system was simulated for 500 time units with randomly generated initial conditions. The initial conditions were drawn from a uniform distribution on [*−*1, 1] for each component *x*_1_, *x*_2_, *x*_3_.

### Additional Methods for Supplementary Figure S2

Each system is simulated with random initial conditions for 2000 time units with the `ode45` integrator in MATLAB. The initial conditions are drawn as in Supplementary Figure S1.

### Additional Methods for Supplementary Figure S3

The return maps for the third dynamical, *x*_3_(*t*), for the simulated network (in black) and the simulated nonlinear dynamical system (in red). The return maps were estimated with 5000 time units worth of data, with the systems integrated with `ode45.`

### Additional Methods for Supplementary Figure S4

The BANFF RNN comprised *N* = 6000 neurons, which were split evenly between *N*_*E*_ = 3000 excitatory and *N*_*I*_ = *N − N*_*E*_ = 3000 inhibitory neurons. The decoders and encoders were randomly generated, with the components of the encoders drawn from the distribution:

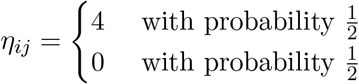

which ensured the encoders are always non-negative. As the encoders were non-negative, the decoders carried the sign of the weight, and therefore the excitatory/inhibitory nature of the presynaptic neuron as ***W*** = ***ηϕ***^*T*^ . The decoders were generated randomly, with the following distribution

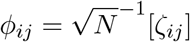

where *ζ*_*ij*_ was a normally distributed random variable and [] was the round operator. With the BANFF RNN weights specified, gradient descent with momentum was used to train the bias currents as in Figures 1-3. The initial learning rate was *ϵ* = 0.02 with the bias currents initialized using the standard normal distribution (mean 0, standard deviation 1). A total of 3 *×* 10^4^ iterations of gradient descent with momentum were used, with a momentum parameter set to 0.9. All dynamical systems and RNNs were integrated with `ode45` under default parameters in MATLAB 2023a (e.g. tolerance, etc.).

### Additional Methods for Supplementary Figure S5

For each dynamical system, the initial point was fed to the network and was a random perturbation from the initial point of the training set. The network was simulated in a closed-loop mode, and the phase portrait was plotted as the first dimension versus the second dimension of the network output (dynamical system).

### Additional Methods for Supplementary Figure S6

For each dynamical system, the initial point was fed to the network and was a random perturbation from the initial point of the training set. The network was simulated in a closed-loop mode, and the return map was plotted as the magnitude of the *n*^th^ peak versus the (*n* + 1)^th^ peak of the third dimension of the network output (dynamical system).

### Notation

- *τ* — membrane (or neuronal) time constant of the recurrent units.
- ***r*** *∈* ℝ^*N ×*1^ — vector of filtered neuronal firing rates.
- ***W*** *∈* ℝ^*N ×N*^ — recurrent synaptic weight matrix.
- ***b*** *∈* ℝ^*N ×*1^ — vector of neuronal bias currents (learnable parameters).
- *f* (*·*) — neuronal transfer function; here *f* (*x*) = tanh(*x*).
- ***ŷ***(*t*) *∈* ℝ^*k×*1^ — decoded output of the network.
- ***ϕ*** *∈* ℝ^*N ×k*^ — readout weight matrix mapping ***r*** to ***ŷ***.
- ***η*** *∈* ℝ^*N ×k*^ — encoder matrix in the low-rank decomposition ***W*** = ***ηϕ***^*T*^ .
- *k* — rank of the recurrent weight matrix ***W***, equal to the dimensionality of the target dynamics.
- *l* — index over target/output dimensions (1 ≤ *l* ≤ *k*).
- *j* — index over neurons (1 ≤ *j* ≤ *N*).
- *n*_*s*_ — number of samples used for discrete loss evaluation.
- *L*(*·, ·*) — general loss or cost function comparing network output to target.
- 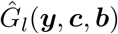— *l*-th component of the network’s predicted dynamics function.
- *G*_*l*_(***y, c***) — *l*-th component of the supervisor (target) dynamics function.
- *f*^*′*^(*·*) — derivative of the transfer function *f* (*·*).
- *ϵ* — learning rate in gradient descent updates.
- *µ* — momentum parameter in gradient descent with momentum.
- ***θ***_*n*_ — momentum term in gradient descent updates at iteration *n*.
- 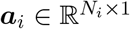 — activation vector of layer *i* in feedforward networks.
- 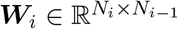 — weight matrix from layer *i −* 1 to *i* in feedforward networks.
- 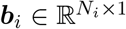 — bias vector for layer *i* in feedforward networks.
- *N*_*i*_ — number of neurons in layer *i*.
- *m*^(*k*)^, *v*^(*k*)^ — first and second moment estimates in Adam optimiser at epoch *k*.
- *α* — learning rate in Adam optimiser.
- *β*_1_, *β*_2_ — exponential decay rates for moment estimates in Adam optimiser.
- *ϵ* (in Adam context) — small constant for numerical stability.
- *L*_MSE_ — mean-squared-error loss.
- *L*_CE_ — cross-entropy loss.
- *y*_*j*_ — *j*-th target output value in supervised learning.
- *ŷ*_*j*_ — *j*-th predicted output value in supervised learning.
- *p*_*j*_ — *j*-th target probability in classification tasks.
- 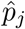 — *j*-th predicted probability in classification tasks.
- 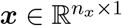 — state vector in control/dynamical system tasks.
- 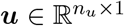 — control input vector.
- ***A, B*** — state-space system matrices in control tasks.
- ***Q, R*** — weighting matrices in quadratic cost function.
- ***K*** — optimal state feedback gain matrix.
- ***P*** — solution to the algebraic Riccati equation.
- *J* — quadratic cost functional in control tasks.
- *t* — continuous time variable.

## Supplementary Information

## Supplementary Appendix 1 Shift of Firing Rates

Here, we will show that BANFF RNNs with negative firing rates, for example with the transfer function *f* (*x*) = tanh(*x*), can be transformed into RNNs with exclusively positive firing rates by redefining the bias-currents.

In particular, 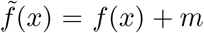 where *m* is some constant such that *f* (*x*) + *m* ≥ 0. Then, consider the RNN equations:

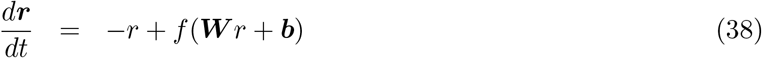

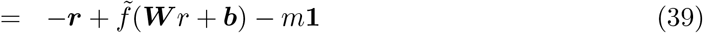

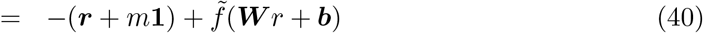

where **1** is an *N ×* 1 vector of 1s in each element. Then, define 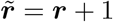:

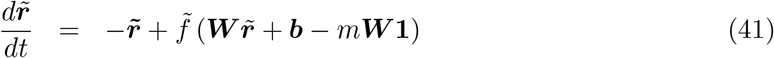

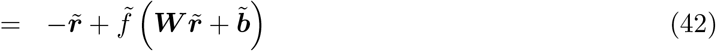

which corresponds to an RNN with all positive firing rates. Note that the weight matrix ***W*** of the original RNN corresponds to the weight matrix of the transformed RNN with strictly positive firing rates, and thus it is sufficient to consider Dale’s Law with the transfer function *f* (*x*) = tanh(*x*) with a change of variables transforming the network into one with positive firing rates, which also respects Dale’s Law.

## Supplementary Appendix 2 Rapid Switching Derivation

Here, we will derive the sufficient conditions under which low-rank BANFF RNNs converge to solutions to the dynamical system 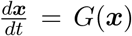 under the assumption of time-varying bias currents.

First, note that the low-dimensional dynamics satisfy the following:

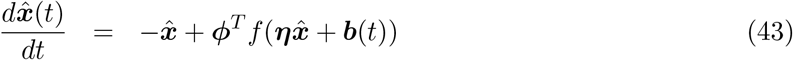

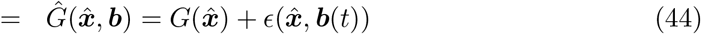

where *ϵ*(***x, b***) denotes the error between the equation governing the target dynamics 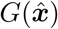 and 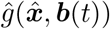. In general, we will assume that ***b***(*t*) will be generated such that the 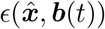 satisfies the following:

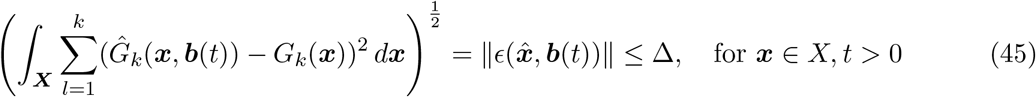

for some Δ *>* 0 where the norm is the conventional *L*_2_ norm over the domain ***X***. One such ***b***(*t*) is given by

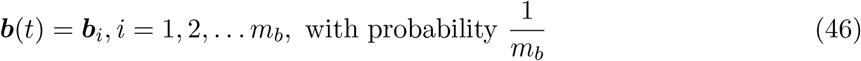

where

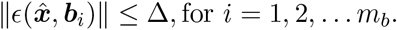

Then, we consider the following:

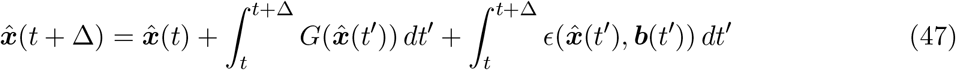

which implies that

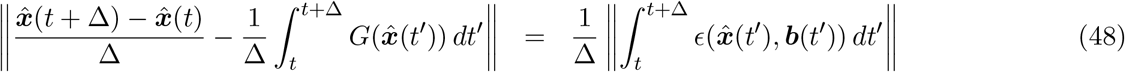

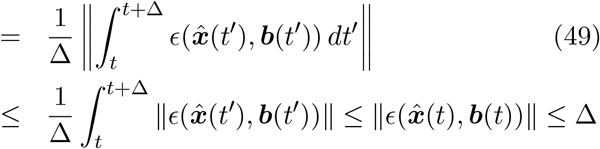

Through the mean-value theorem for integrals, and the fact that 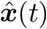 is continuous, we know that the left hand side of equation (48) converges to a solution to the differential equation given by ***x***^*′*^(*t*) = *G*(***x***) as Δ → 0. When considering the switching bias current given by equation, (46), this derivation implies that if the *L*_2_ error for each bias current is bounded by common bound Δ, then the approximant 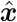 approximates ***x***(*t*) to at least *O*(Δ).

## Supplementary Appendix 3 Feedforward Neural Network Tasks

### Classification Tasks

For all classification tasks, pre-existing labelled datasets were used.

#### Iris Species task

The task was to classify the species of an iris from four features: the sepal length, sepal width, petal length and petal width. Three species were present in the dataset: Setosa, Versicolor and Virginica.

#### Breast Cancer task

The task was to classify a breast mass as either malignant or benign, given details from an image of a fine needle aspirate of a breast mass. The mean, standard error and mean of the three largest values for ten measurements of the cell nucleus were used as 30 features. The ten measurements were: radius, texture, perimeter, area, smoothness, compactness, concavity, number of concave points, symmetry and fractal dimension.

#### Car Quality task

the task was to classify a car either ‘unacceptable’, ‘acceptable’, ‘good’ or ‘very good’ when provided with the following six features for the car: purchase price, maintenance price, number of doors, capacity (number of people), size of the boot (luggage capacity) and the estimated safety.

#### Mushroom task

the task was the classify a mushroom as either poisonous or edible given the following 22 features for each mushroom: cap shape, cap surface, cap colour, bruises, odour, gill attachment, gill spacing, gill size, gill colour, stalk shape, stalk root, stalk surface above ring, stalk surface below ring, stalk colour above ring, stalk colour below ring, veil type, veil colour, ring number, ring type, spore print colour, population and habitat.

#### MNIST-type tasks

In this group, the task was to classify a 28 *times*28 pixel input image as one of ten classes. For MNIST, all Afro-MNIST and Kuzushiji-MNIST tasks, the images were handwritten digits, and the classes were those digits (e.g. the digits 0 to 9). For the Fashion-MNIST task, the input images were items of clothing belonging to one of the following ten categories: T-shirt/top, trousers, jumper, dress, coat, sandals, shirt, trainers, bag or ankle boot.

### Regression Tasks

For all classification tasks, pre-existing labelled datasets were used.

#### Abalone Age task

The task was to output the age of an abalone given the following eight features: sex, length, diameter perpendicular to length, height, weight, shucked weight, viscera weight and shell weight.

#### Car Price task

The task was to output the price of a used car in the United Kingdom (all Toyota), given the following eight features: registration year, transmission, mileage, fuel type, tax, miles per gallon and engine size.

### Dynamical Systems Tasks

**Supplementary Table 1.**
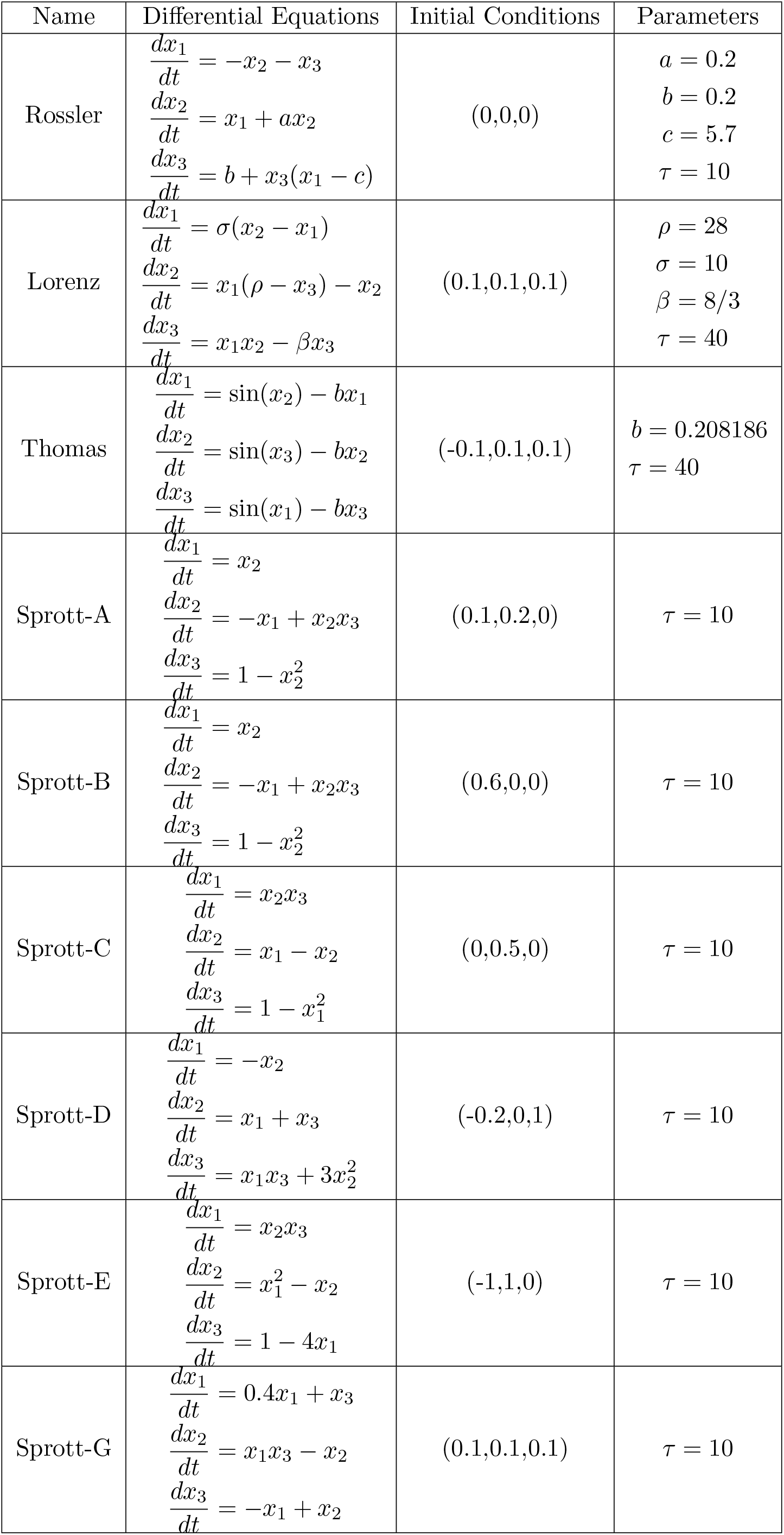
List of Dynamical Supervisors 1.

**Supplementary Table 2.** List of Dynamical Supervisors 2.

| Name | Differential Equations | Initial Conditions | Parameters |
| --- | --- | --- | --- |
| Sprott-I | $\frac{dx_1}{dt} = -0.2x_2$ $\frac{dx_2}{dt} = x_1 + x_3$ $\frac{dx_3}{dt} = x_1 + x_2^2 - x_3$ | (0,0.3,0) | $\tau = 10$ |
| Sprott-J | $\frac{dx_1}{dt} = 2x_3$ $\frac{dx_2}{dt} = -2x_2 + x_3$ $\frac{dx_3}{dt} = -x_1 + x_2^2 + x_2$ | (-0.1,1,0.1) | $\tau = 10$ |
| Sprott-K | $\frac{dx_1}{dt} = x_1x_2 - x_3$ $\frac{dx_2}{dt} = x_1 - x_2$ $\frac{dx_3}{dt} = x_1 + 0.3x_3$ | (1,1,2) | $\tau = 10$ |
| Sprott-M | $\frac{dx_1}{dt} = -x_3$ $\frac{dx_2}{dt} = -x_1^2 - x_2$ $\frac{dx_3}{dt} = 1.7 + 1.7x_1 + x_2$ | (1,0.8,0) | $\tau = 10$ |
| Sprott-N | $\frac{dx_1}{dt} = -2x_2$ $\frac{dx_2}{dt} = x_1 + x_3^2$ $\frac{dx_3}{dt} = 1 + x_2 - 2x_3$ | (4.5,1,0) | $\tau = 10$ |
| Sprott-O | $\frac{dx_1}{dt} = x_2$ $\frac{dx_2}{dt} = x_1 - x_3$ $\frac{dx_3}{dt} = x_1 + x_1x_3 + 2.7x_2$ | (0,0,0.5) | $\tau = 10$ |
| Sprott-Q | $\frac{dx_1}{dt} = -x_3$ $\frac{dx_2}{dt} = x_1 - x_2$ $\frac{dx_3}{dt} = 3.1x_1 + x_2^2 + 0.5x_3$ | (1,0,0) | $\tau = 10$ |
| Sprott-S | $\frac{dx_1}{dt} = -x_1 - 4x_2$ $\frac{dx_2}{dt} = x_1 + x_3^2$ $\frac{dx_3}{dt} = 1 + x_1$ | (0,0,1) | $\tau = 10$ |
| Van der Pol | $\frac{dx_1}{dt} = \mu(x_1 - \frac{x_1^3}{3} - x_2)$ $\frac{dx_2}{dt} = \frac{x_1}{\mu}$ | (0.2,0.3) | $\mu = 5$<br>$\tau = 10$ |

**Supplementary Table 3.**
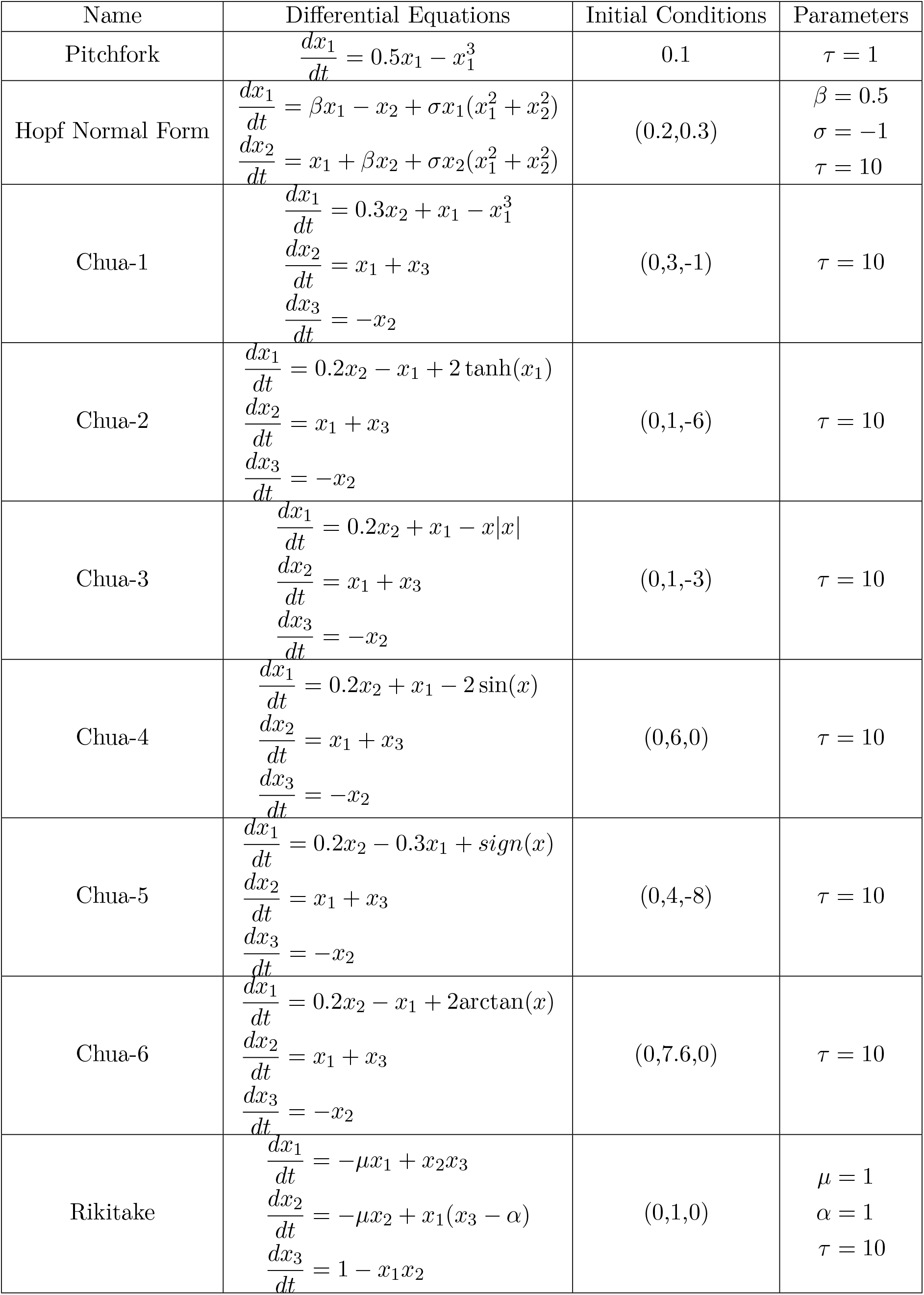
List of Dynamical Supervisors 3.

**Supplementary Table 4.**
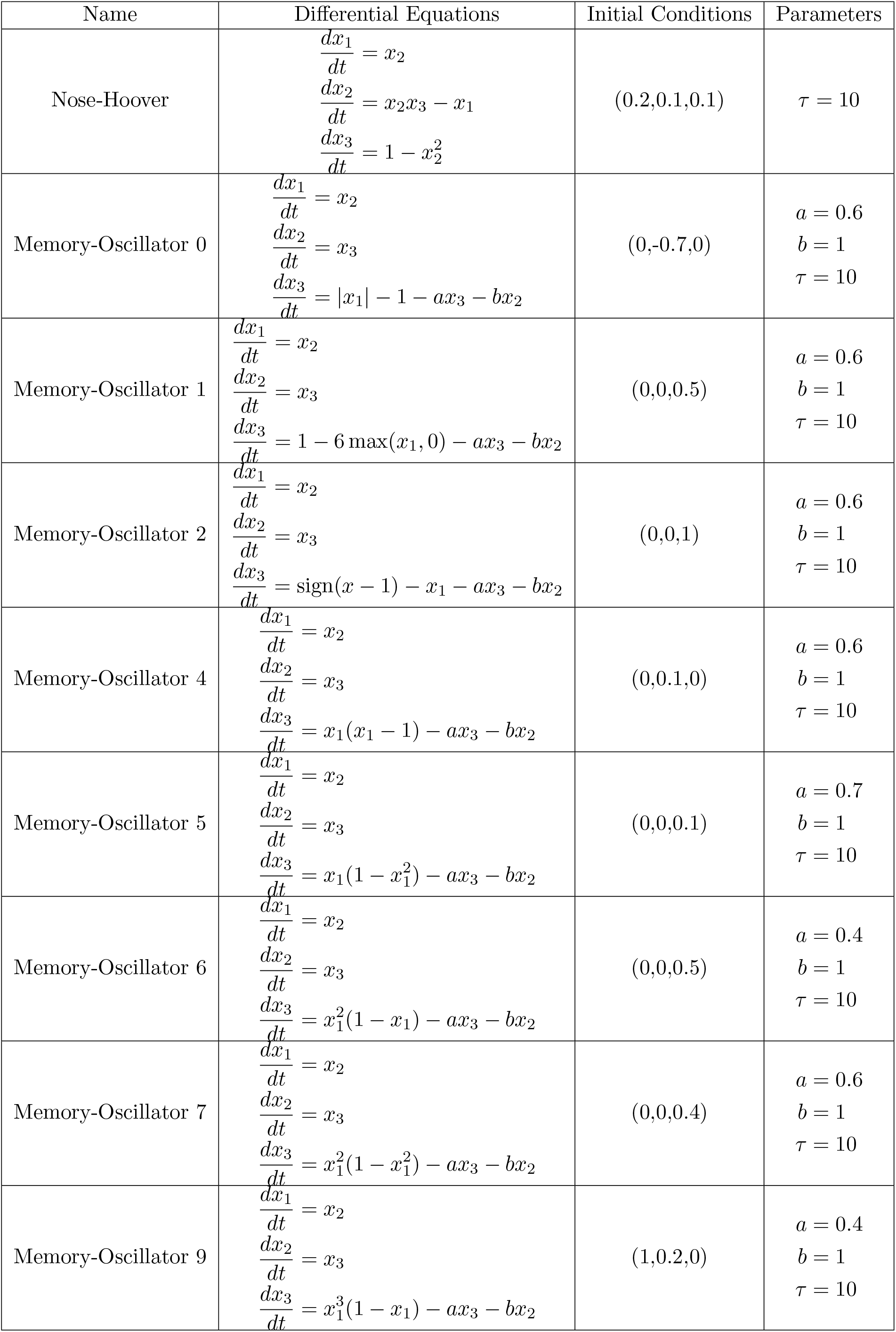
List of Dynamical Supervisors 4.

**Supplementary Table 5.** List of Dynamical Supervisors 5.

| Name | Differential Equations | Initial Conditions | Parameters |
| --- | --- | --- | --- |
| Memory-Oscillator 13 | $\frac{dx_1}{dt} = x_2$ $\frac{dx_2}{dt} = x_3$ $\frac{dx_3}{dt} = 6 \tanh(x_1) - 3x_1 - ax_3 - bx_2$ | (0,0,1) | $a = 1$<br>$b = 1$<br>$\tau = 10$ |
| Memory-Oscillator 14 | $\frac{dx_1}{dt} = x_2$ $\frac{dx_2}{dt} = x_3$ $\frac{dx_3}{dt} = 6 \arctan(x_1) - x_1 - ax_3 - bx_2$ | (0,1,6) | $a = 0.6$<br>$b = 1$<br>$\tau = 10$ |
| Memory-Oscillator 15 | $\frac{dx_1}{dt} = x_2$ $\frac{dx_2}{dt} = x_3$ $\frac{dx_3}{dt} = x_1 - 0.5 \sinh(x_1) - ax_3 - bx_2$ | (0,0,1) | $a = 0.6$<br>$b = 1$<br>$\tau = 10$ |

## Supplementary Figures

**Supplementary Figure S1.**
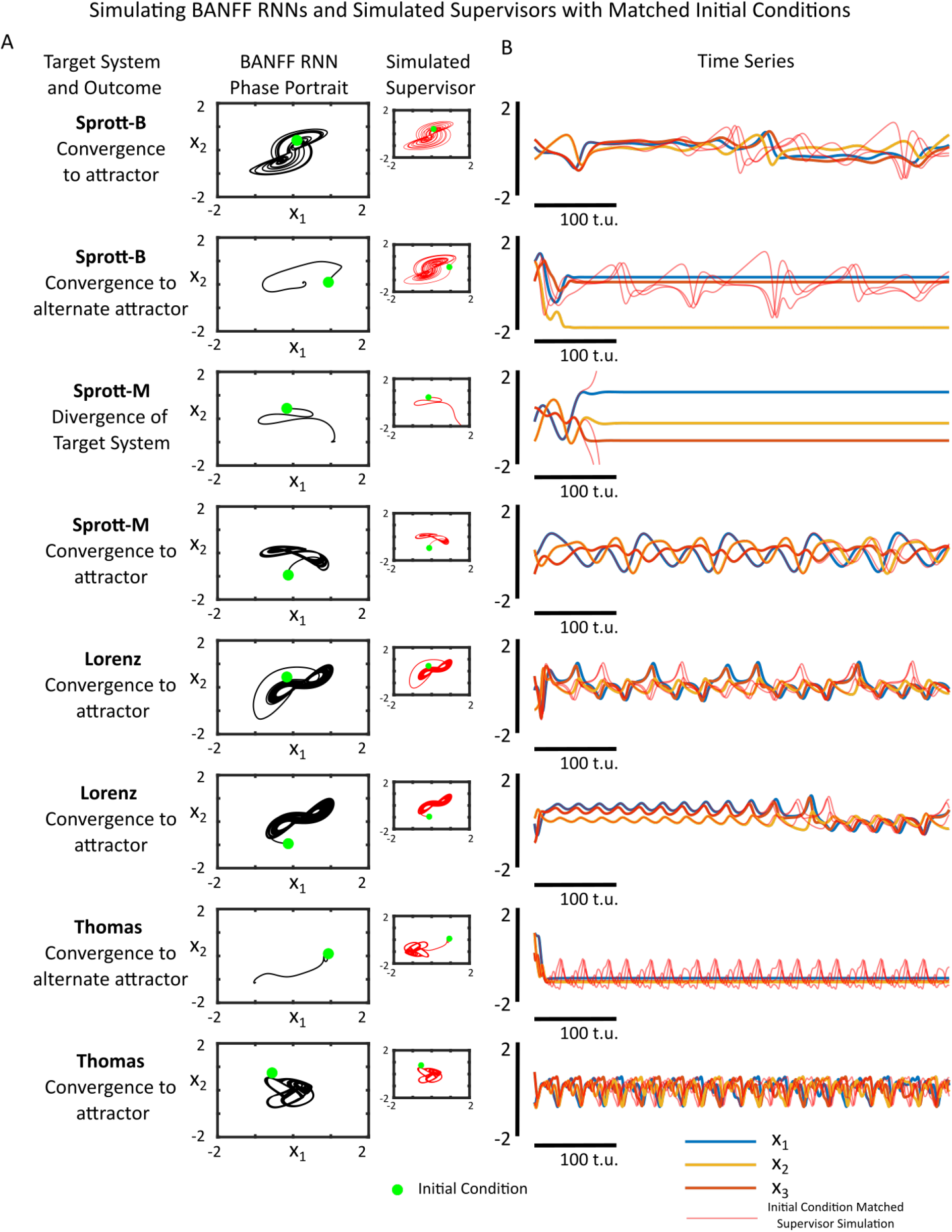
Simulating Trained BANFF RNNs with Different Initial Conditions. **(A)** The phase portrait for a single BANFF RNN trained on 4 separate target supervisors with two different initial conditions each. The BANFF RNN was simulated for 500 time units, with different initial conditions, each drawn from a uniform distribution on [*−*1, 1]^3^. The simulated supervisor with an identical initial condition is shown as an inset. With some initial conditions, the BANFF RNN converges to the target attractor, while with other initial conditions, the BANFF RNN can converge to alternate attractors (e.g. fixed points) **(B)** The time series for the simulated BANFF RNNs (blue/orange/yellow lines) and the simulated supervisor with matched initial conditions (red lines).

**Supplementary Figure S2.**
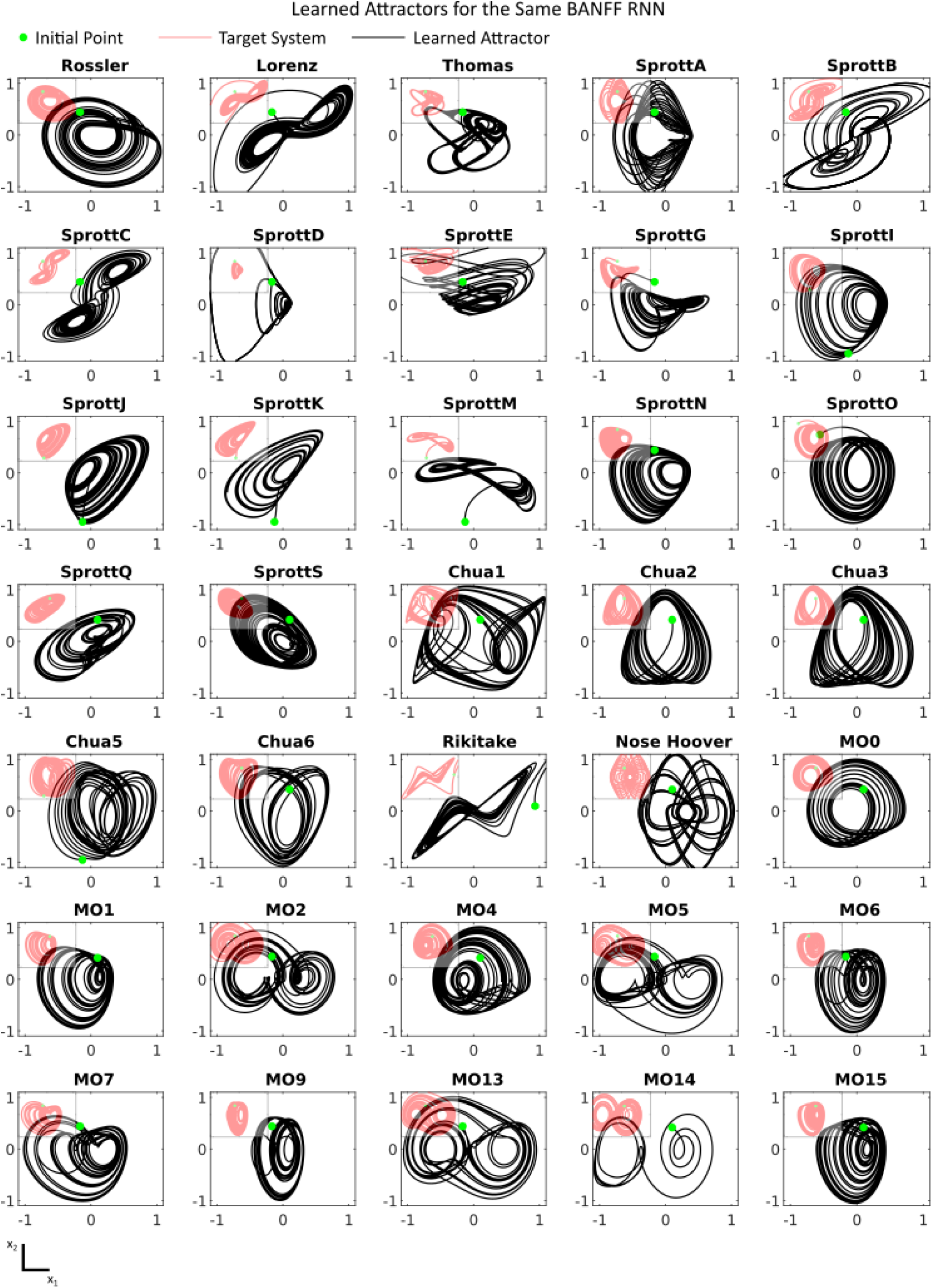
The Same BANFF RNN Mimicking 35 Different Chaotic Systems with Learned Bias Currents: The Phase portrait for a single BANFF RNN consisting of 6000 neurons with fixed weights and different bias currents. The applied bias currents were trained with gradient descent so that the BANFF RNN could reproduce the chaotic attractors of the corresponding nonlinear dynamical system (red inset). In most cases, the BANFF RNN successfully reproduces the attractor, while in some cases, a heavily folded limit cycle that is visually similar to the expected chaotic attractor (e.g. MO14) is produced by the RNN instead.

**Supplementary Figure S3.**
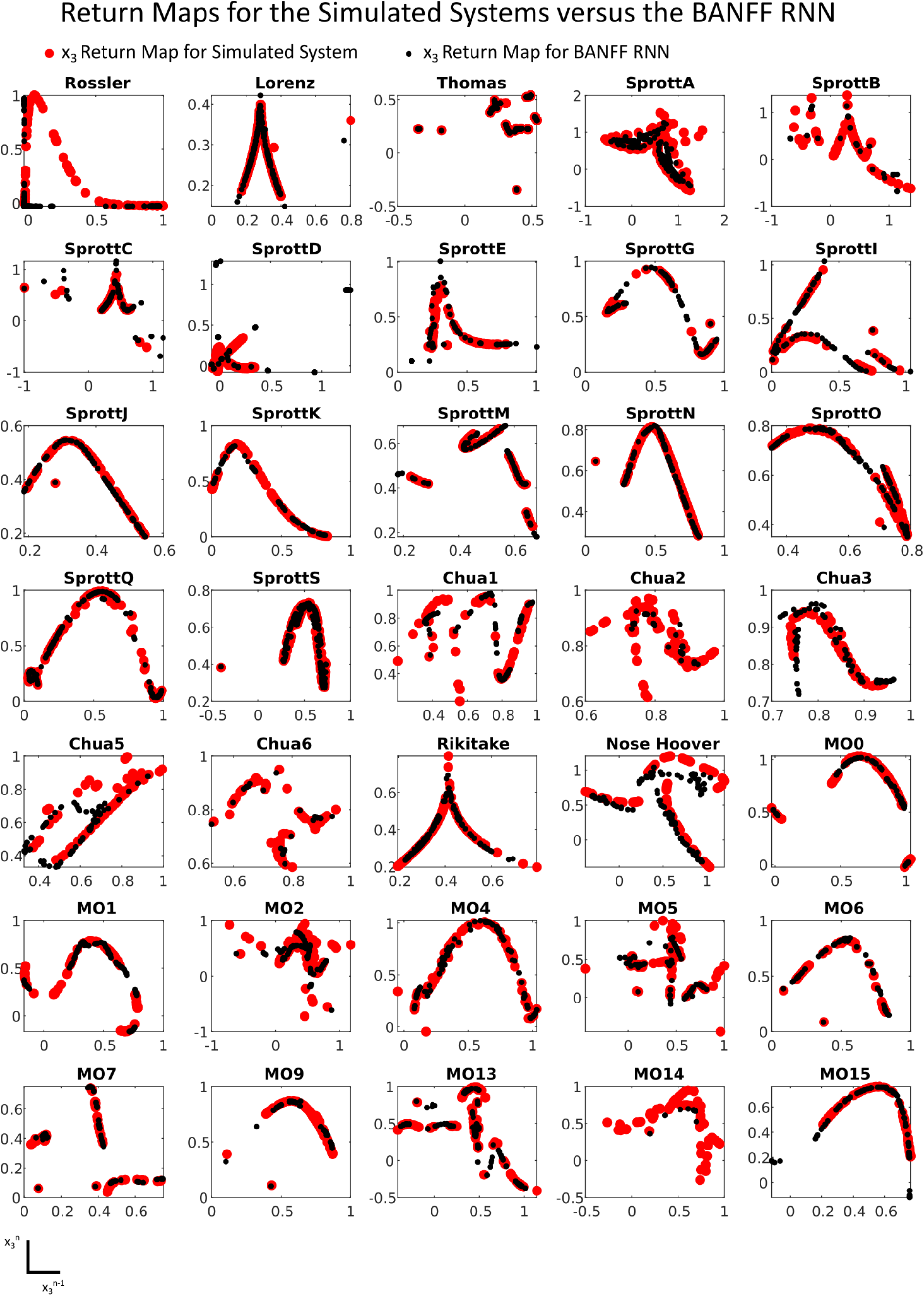
The Same BANFF RNN Approximating the Return Maps of 35 Separate Dynamical Systems: The return maps for simulated nonlinear dynamical systems (in red) versus a single trained BANFF RNN for 35 separate dynamical systems. The return map is defined as the *n −* 1th peak of *x*_3_(*t*) versus the *n*th peak of *x*_3_(*t*). Accurate mimicry of the return map indicates accurate mimicry of the local dynamics of the chaotic systems.

**Supplementary Figure S4.**
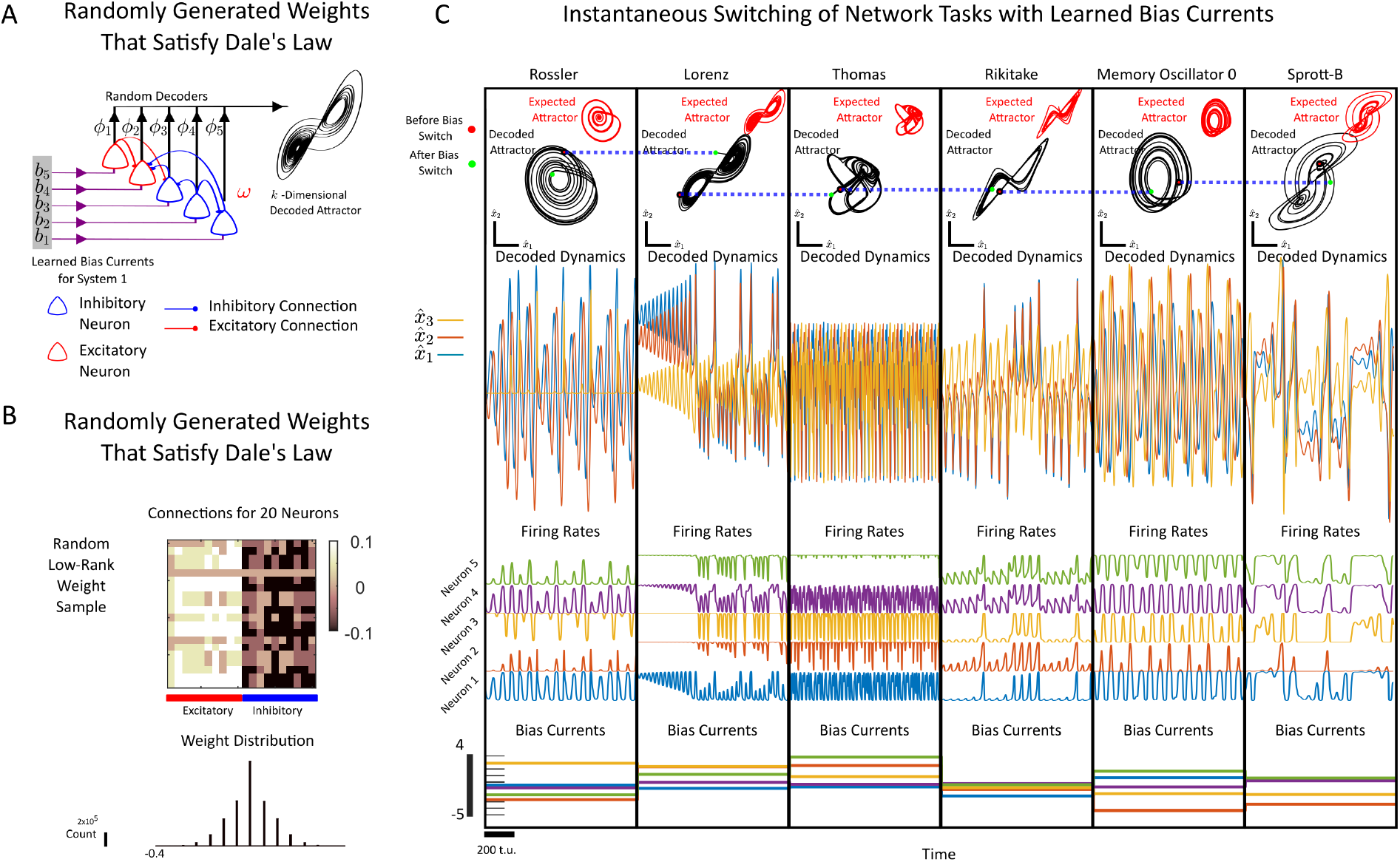
The Bias Adaptive Neural Firing Framework applied to networks with weights that satisfy Dale’s Law. **(A)** Schematic: A low-rank RNN with fixed weights is trained to decode out the dynamics of different supervisors by changing the input bias currents to the neurons. The network is composed of inhibitory and excitatory connections only, which respect the presynaptic neuron’s designation as an inhibitory/excitatory neuron. Inhibitory neurons only project negative (inhibitory) weights, while excitatory neurons only project positive (excitatory) weights. **(B)** (Top) Randomly generated low-rank weights that respect Dale’s Law. The connections shown are for 10 randomly selected neurons, with the first 10 indices corresponding to excitatory neurons and the next 10 indices corresponding to inhibitory neurons. The heatmap denotes the excitatory/inhibitory strength. The distribution of weights is shown below the weight matrix, with the weights also having discrete values. **(C)** (First Row) The phase portrait of the decoded attractor (black) versus the expected attractor (red) for 6 different nonlinear dynamical systems. The red /green dots denote the state of the network before a bias switch (red) and after a bias switch (green). (Second Row) The time series for the decoded dynamics after each switch for 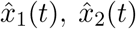, and 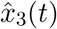. (Third Row) The firing rates *r*_*i*_(*t*) for 5 separate neurons in the BANFF RNN. (Fourth Row) The bias currents applied to the BANFF RNN. Each bias current is applied for 1000 time units before a switch.

**Supplementary Figure S5.**
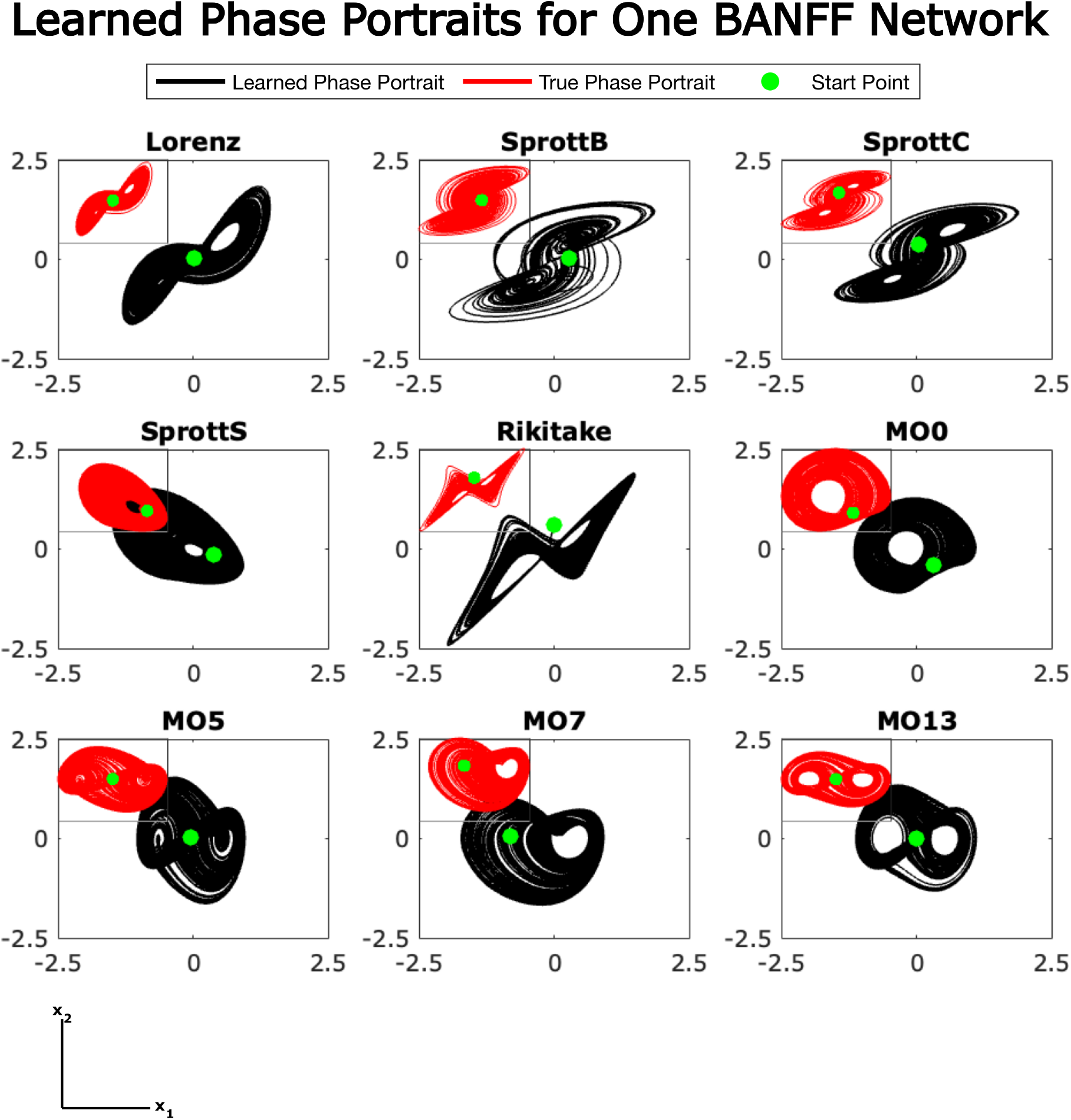
The Phase Portraits for one feedforward BANFF RNN operating in a closed-loop configuration. The phase portraits of the feedforward BANFF network output and the deterministically simulated system for each of the dynamical systems considered. In each case, the plot is the first dimension of the dynamical system *x*_1_ with respect to the second dimension *x*_2_.

**Supplementary Figure S6.**
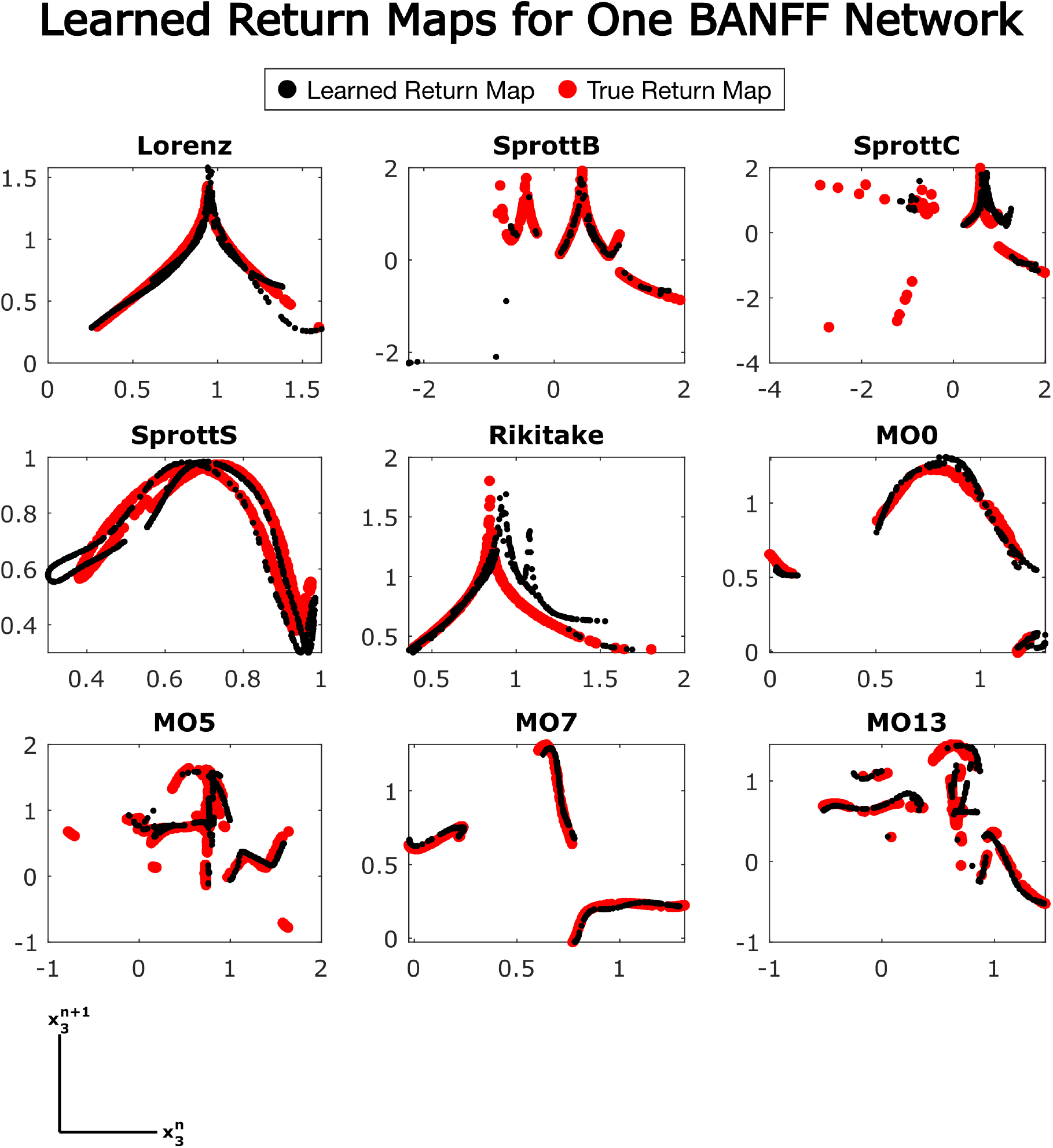
Return Maps for a Feedforward BANFF RNN Operating in Closed Loop. The return maps of the feedforward BANFF network output and the deterministically simulated system for each of the dynamical systems considered. In each case, the plot is the amplitude of the *n*^th^ peak with respect to the amplitude of the (*n* + 1)^th^ peak of the third dimension *x*_3_(*t*) of the dynamical system.

